# Anomalous Perception of Biological Motion in Autism: A Conceptual Review and Meta-Analysis

**DOI:** 10.1101/530030

**Authors:** Alessandra Federici, Valentina Parma, Michele Vicovaro, Luca Radassao, Luca Casartelli, Luca Ronconi

**Author notes:** Correspondence for this article should be sent to: Luca Ronconi, PhD; Faculty of Psychology, Vita-Salute San Raffaele University, Milan, Italy, Luca Casartelli, PhD; Child Psychopathology Unit, Scientific Institute IRCCS E. Medea, Bosisio Parini, Lecco, Italy. co-first authorship (Alessandra Federici, Valentina Parma). co-last authorship (Luca Casartelli, Luca Ronconi).

## Abstract

The construct of biological motion (BM) has gained increasing popularity in the last decades and has been widely investigated as a marker of social difficulties in individuals with autism spectrum disorder (ASD). However, studies testing BM have extremely heterogeneous experimental protocols and, moreover, evidence of BM perception anomalies in ASD have been mixed. In this article, we present a meta-analysis investigating the putative anomalies of BM perception in ASD. Through a systematic literature search, we found 27 studies that investigated BM perception in both ASD and typical developing (TD) peers by using point-light display stimuli. A general meta-analysis including all these studies showed a moderate deficit of individuals with ASD in BM processing, but also a high heterogeneity. This heterogeneity was explored in different additional meta-analyses where studies were grouped according to different levels of complexity of the BM task employed (*first-order*, *direct* and *instrumental*), and according to the manipulation of low-level perceptual features (*spatial* vs. *temporal*) of the control stimuli used as comparison. Results suggest that the most severe deficit in ASD is evident when perception of BM is serving a secondary purpose (e.g. inferring intentionality/action/emotion) and, interestingly, that temporal dynamics of stimuli is an important factor in determining BM processing anomalies in ASD. Our results question the traditional understanding of BM anomalies in ASD as a monolithic deficit and suggest a paradigm shift that deconstructs BM into distinct levels of processing and specific spatio-temporal subcomponents.

## 1. Introduction

Our brain is constantly facing a plethora of sensory stimuli that needs to be properly sampled and organized to construct a meaningful perceptual experience. Since seminal studies on point-light displays (PLD) investigating the perception of distinct types of motion patterns such as walking, running and dancing (Johansson, 1973), the robust tuning of the human visual system to biological motion (BM) has represented an intriguing challenge for scientists (Neri, Morrone, & Burr, 1998; Pavlova, 2012; Proffitt, Bertenthal, & Roberts, 1984; Simion, Regolin, & Bulf, 2008). Compared to full body motion, stimuli created with point-light displays permit to profitably separate motion processing from other features such as color or shape, representing an essential tool to investigate the principles behind our perceptual tuning to BM. In light of its implications for cognitive science, developmental and clinical neuroscience, the study of BM processing raised considerable attention in recent years (Rosa Salva, Mayer, & Vallortigara, 2015; J. Thompson & Parasuraman, 2012). In particular, its putative implications in social cognition have been widely debated in the context of Autism Spectrum Disorder (ASD), raising the hypothesis that anomalies in BM processing could be considered a marker or an intermediate phenotype of ASD (Pavlova, 2012). The present work has multiple aims. First, to test whether (and eventually to what extent) the processing of BM – in its most low-level form, namely PLD stimuli - is effectively anomalous in ASD. Second, to consider a different theoretical and conceptual model that deconstructs BM into distinct levels of processing and according to distinct perceptual manipulations. Third, to present a quantitative meta-analysis of putative BM processing anomalies in ASD that follows this alternative conceptualization.

Being characterized by early onset, lifelong and debilitating behavioral symptomatology, ASD represents a serious concern for families, clinicians and generally for societies (Buescher, Cidav, Knapp, & Mandell, 2014; Lord, Elsabbagh, Baird, & Veenstra-Vanderweele, 2018). To date, the clinical diagnosis of ASD is based on behavioral symptoms and according to the DSM 5 it is characterized by restricted and repetitive patterns of behaviors, interests or activities, and difficulties in the social communication/interaction domain (American Psychiatric Association, 2013). Hyper- or hypo-reactivity to sensory stimuli or unusual interest for specific sensory aspects are also considered important features of the disorder (Robertson & Baron-Cohen, 2017a). One of the major concerns in dealing with ASD is the heterogeneity in severity and behavioral manifestations and challenges in its early detection when behavioral signs are less evident (Lai, Lombardo, & Baron-Cohen, 2014). These aspects significantly impact on the clinicians’ ability to provide early and specific interventions (Constantino & Charman, 2016). Although the growing body of studies targeting neural structures implicated in the pathophysiology of ASD has improved our insights on the condition (Fatemi et al., 2012; Kucharsky Hiess et al., 2015; Paul, Corsello, Kennedy, & Adolphs, 2014; Wang, Kloth, & Badura, 2014), the neurobiology of ASD is far from being clarified (Di Martino et al., 2014; Ecker & Murphy, 2014; Haar, Berman, Behrmann, & Dinstein, 2016; Lai et al., 2014; Stoner et al., 2014). Similarly, neither brain functional and structural connectivity approaches (Khan et al., 2015; J. D. Lewis et al., 2017) nor genetics (Geschwind, 2011; Richards, Jones, Groves, Moss, & Oliver, 2015) have definitely solved the puzzle. Efforts in understanding ASD are hampered by substantial disconnection between neurobiological findings aiming to characterize the neural/genetic components associated with the condition, and its core behavioral phenotype. Thus, research investigating endophenotypes of ASD, i.e. measurable and quantifiable links between neurobiological underpinnings and behavioral symptoms (Gottesman & Gould, 2003; Kendler & Neale, 2010), represents a critical challenge for the future (Homberg et al., 2016; Klin, Shultz, & Jones, 2015). In recent years, a growing body of studies on endophenotypes of ASD has been performed exploring complex functions such as visual attention (Gammer et al., 2015; Keehn, Müller, & Townsend, 2013; Robertson, Kravitz, Freyberg, Baron-Cohen, & Baker, 2013; Ronconi, Devita, Molteni, Gori, & Facoetti, 2018; Ronconi, Gori, et al., 2018; Ronconi, Gori, Ruffino, Molteni, & Facoetti, 2013), visual perception (Ewing, Pellicano, & Rhodes, 2013; Foss-Feig, Tadin, Schauder, & Cascio, 2013; Kröger et al., 2014; Vlamings, Jonkman, van Daalen, van der Gaag, & Kemner, 2010), motor cognition (Ansuini, Podda, Battaglia, Veneselli, & Becchio, 2018; Becchio, Pierno, Mari, Lusher, & Castiello, 2007; Casartelli, Molteni, & Ronconi, 2016; Cattaneo et al., 2007; D’Angelo & Casali, 2012; Rochat et al., 2013) and sensory processing (Collignon et al., 2013; Endevelt-Shapira et al., 2018; Kwakye, Foss-Feig, Cascio, Stone, & Wallace, 2011a; Parma, Bulgheroni, Tirindelli, & Castiello, 2013; Robertson & Baron-Cohen, 2017a; Rozenkrantz et al., 2015).

A significant interest in the literature has been directed to the idea that individuals with ASD have a different perceptual experience of the world and, even more critical for our aims, that sensory/perceptual anomalies in ASD are not just a secondary effect of reduced social interactions. Accordingly, sensory/perceptual anomalies would represent a key and primary component both in terms of symptom development and pathophysiology (Robertson & Baron-Cohen, 2017a). A specific focus has been oriented to BM processing in ASD. Following the first report of BM impairment in ASD (Moore, Hobson, & Lee, 1997), the number of studies investigating this question has constantly increased over the years (see Figure 1). However, factors such as heterogeneous experimental designs, weak understanding of potential confounds and inconsistency among the classes of information conveyed by BM may rise various misunderstandings. To date, there is a general and quite vague agreement on the supposed BM anomalies in ASD, which however lacks both a clear epistemological awareness of the construct of BM and a quantitative evaluation of these alleged anomalies (Pavlova, 2012; Pelphrey & Carter, 2008; Redcay, 2008).

**Figure 1.**
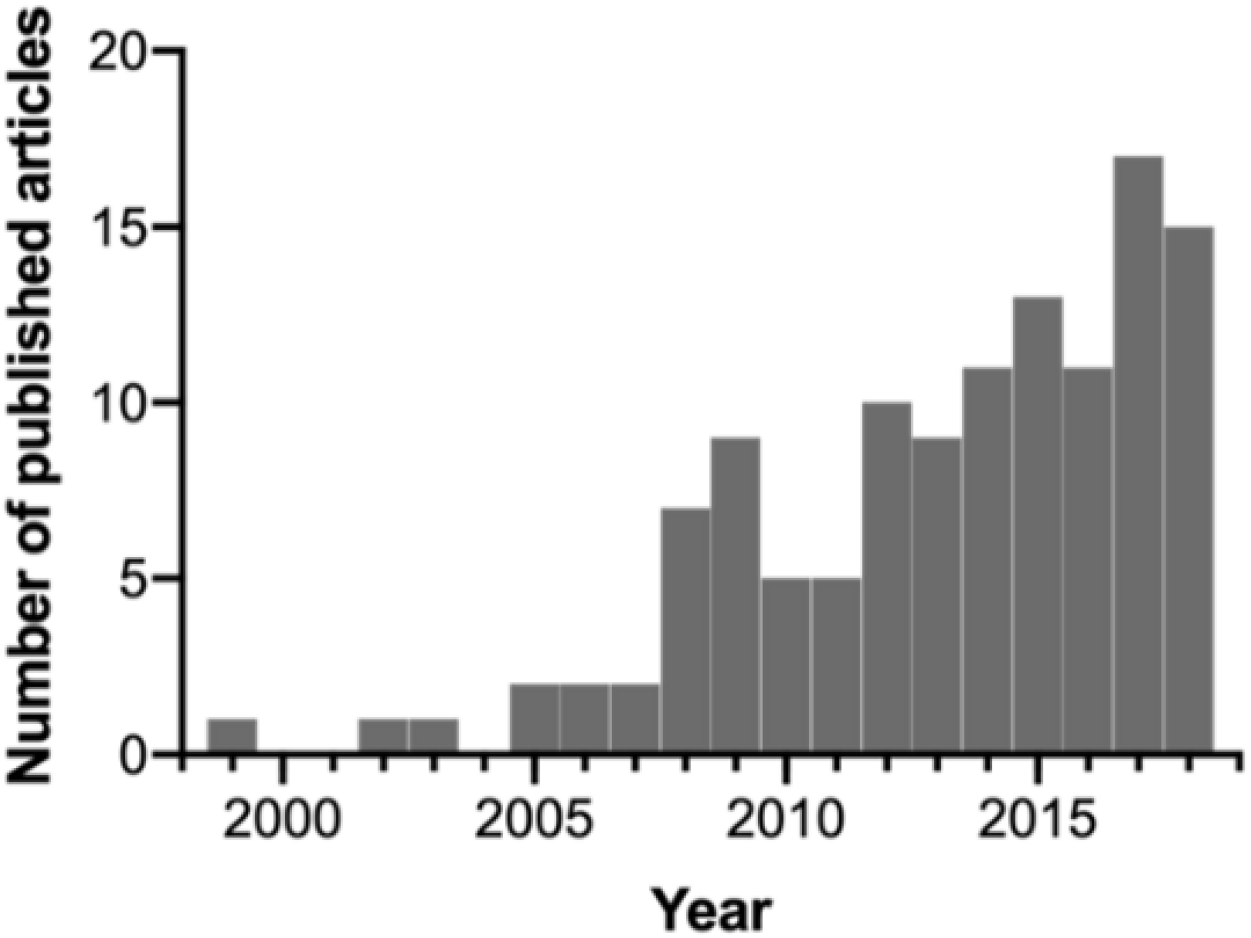
Number of articles published per year as indexed in PubMed with the general search using the keywords “biological motion autism”.

### 1.1 Biological motion processing: an early emerging ability

We may expect that a phylogenetically highly preserved mechanism, such as BM processing, also shows an early developmental progression during the ontogenesis. However, it still remains to be clarified whether, and eventually to what extent, the tuning to BM is dependent on experience. A seminal study in infancy indicates that at 3 months of age infants are able to discriminate BM stimuli vs. their scrambled and upright counterparts (Bertenthal, Proffitt, & Cutting, 1984). This trend is observed also at 4-5 months of age (Bertenthal et al., 1984; Fox & McDaniel, 1982). More recently, a convincing argument in favor of the hypothesis that BM processing is largely independent from experience was provided by a study testing the ability of newborns to discriminate BM stimuli (Simion et al., 2008). The sensitivity to BM was tested in 2-day-old babies using hen-walking animations as compared to nonbiological stimuli (i.e., the same pattern of elements moving in a random manner) and inverted (upside -down) hen-walking stimuli. The use of hen stimuli allowed to exclude any possible (although remote) learning mechanisms taking place in the first 2 days of life. The results indicate that newborns manifest a spontaneous preference for BM stimuli, supporting the idea that this processing is largely (or even completely) independent from experience (Simion et al., 2008). Although the authors could not rule out the possible role of fetal experience in shaping BM predispositions (see for example Pavlova, 2012), these findings supported a certain degree of innate or nearly innate tuning of the human perceptual system to BM.

The developmental trajectory of the sensitivity to BM stimuli has also been extensively explored in comparison to global motion (Hadad, Maurer, & Lewis, 2011). Testing different groups of children (aged 6-8, 9-11 and 12-14 years, respectively) and adults, Hadad and colleagues (2011) claimed that developmental trajectories for coherent motion and BM are similar. In both cases, the authors found that thresholds achieved adult-like profiles around the age of 14, following a sort of monotonic developmental trajectory of improvement from childhood. In contrast, testing before surgery the BM processing and global motion in patients that suffered from early visual deprivation because of dense bilateral congenital cataracts, showed that marked deficits in global motion are not accompanied with similar difficulties in BM processing (Hadad, Maurer, & Lewis, 2012). Importantly, a recent EEG study showed that the BM perception is not only preserved at the behavioral level, but also that the same neural processing is involved. Indeed, in patients with congenital cataracts, the N1 component of event-related potentials was modulated post-surgery by the processing of BM as in the control group (Bottari et al., 2015). Spared abilities in the processing of BM after early visual deprivation are surprising in light of the data reporting impairments in similar cohorts of patients both in global motion (T. L. Lewis et al., 2002) and in holistic face processing (Le Grand, Mondloch, Maurer, & Brent, 2004; Robbins, Nishimura, Mondloch, Lewis, & Maurer, 2010).

Taken together, these data indicate a composite picture in which clear similarities (Hadad et al., 2011) and marked differences (Hadad et al., 2012) between basic and more complex mechanisms (e.g., global motion and BM processing) coexist (see also for example Mather, Battaglini, & Campana, 2016). Such coexistence should stimulate further studies aiming at characterizing shared developmental patterns vs. neuroplastic mechanisms. For shared patterns, a classical approach refers to the well-established neurocostructivist view supporting the idea that very early, not-detailed, domain-general functions impact on the development of more complex, domain-specific, and even “social” functions (Karmiloff-Smith, 1998; Ronconi, Molteni, & Casartelli, 2016). For compensatory mechanisms, examples from heterogeneous clinical pictures may provide intriguing insights (Hadad et al., 2012; for example, related to other functions, see also Casartelli & Parma, 2017; Rezlescu, Barton, Pitcher, & Duchaine, 2014). This overview of both BM phylogenesis and ontogenesis suggests that oversimplifying this construct, as if it was to convey a monolithic process, can be misleading and that a paradigm shift in approaching BM is needed.

### 1.2 Motor, spatio-temporal and social aspects of biological motion processing

It has been hypothesized that the ability to detect, recognize and interpret BM is related, in different ways, to the expertise in performing similar movements or actions and, to some extent, also to one’s social abilities (Pavlova, 2012). Beyond the attractiveness of this view, whether and how the ability to process BM is linked *tout court* to the expertise in producing similar movements or actions requires a cautious interpretation. Recent experimental and theoretical investigations on mirror mechanisms clearly indicate that humans benefit from their own motor representations to understand the actions of others directly (“motorically”, “from the inside”; Rizzolatti & Sinigaglia, 2016). Notably, mirroring phenomena may occur at distinct level of abstractness, as if we are able to represent distinct components of an action in a motor way (Rizzolatti, Fogassi, & Gallese, 2001; Rizzolatti & Sinigaglia, 2016), and they have been hypothesized to be anomalous in ASD (Casartelli et al., 2016; Trevarthen & Delafield-Butt, 2013).

Two studies reporting seemingly diverging results may help in furnishing further insights on this point. First, adolescents born preterm and suffering from periventricular leukomalacia (PVL), a clinical condition characterized by white matter lesions near the lateral ventricles, were tested in their ability to process BM (Krägeloh-Mann et al., 1999; Pavlova, 2012; Pavlova, Staudt, Sokolov, Birbaumer, & Krägeloh-Mann, 2003). These participants varied in their motor ability, ranging from complete walking disability to typical or near-typical motor patterns (Pavlova et al., 2003). Impairment in the execution of actions with the lower and upper limbs correlated with the lateral extent of periventricular lesions but did not correlate with the performance at the BM task. Interestingly, sensitivity to the processing of BM negatively correlated with the volume of the lesions over the parieto-occipital complex, whereas no correlation over the frontal and temporal regions was found (Pavlova et al., 2003). Second, the link between action perception and action execution was explored by Casile and Giese (2006), who tested whether the acquisition of novel motor behaviors could improve the processing of BM stimuli. The authors demonstrated that nonvisual motor training selectively improves the visual recognition of novel BM patterns, thus suggesting that motor learning, independently of visual feedback given that participants were blindfolded, impacts on BM recognition (Casile & Giese, 2006). Taken together, these two studies outline a more articulated picture. The hypothesis that BM recognition is linked to some extent to the abilities in producing specific motor patterns is fascinating, but its simplistic interpretation should be overcome (Casile & Giese, 2006). At the same time, BM processing seems to benefit from more efficient and unexpected compensatory mechanisms, in turn suggesting that functional networks supporting BM processing are more complex (Pavlova et al., 2003 see also Sokolov, Erb, Grodd, & Pavlova, 2014).

Another source of debate in the literature is how spatio-temporal features of BM stimuli impact on BM processing itself. BM processing entails both spatial and temporal dynamics, but the exact contribution of these components and individual tolerance to their perturbation have not been clarified in ASD (Hadad et al., 2011; Troje & Westhoff, 2006). This may offer insights toward the understanding of apparently controversial findings reported in the ASD literature, given that being tolerant to spatial and/or temporal perturbations are supported by potentially distinct neural mechanisms. Recent evidence refined traditional views about timing, attributing also to the cerebellum a key role in it (Bareš et al., 2018). Nowadays, cerebellar contribution to non-motor functions such as sensory and perceptual ones are well-established (Baumann et al., 2015; Buckner, 2013). Interestingly, also the contribution of the cerebellum in the pathophysiology of ASD (Fatemi et al., 2012) and – more specifically – in ASD sensory/perceptual anomalies have been recently explored (Casartelli, Riva, Villa, & Borgatti, 2018; Wang et al., 2014).

The complexity of networks supporting BM processing and neuroplasticity mechanisms that compensate, at least partially, the recognition of BM may be considered cues of the functional relevance of BM. A classical view suggests that the perception of BM is tightly linked with individual abilities in the social domain, leading back this hypothesis to the evolutionary well-established benefit in distinguishing efficiently biological from nonbiological motion. This “social” interpretation of BM posits that performance in BM tasks may serve as benchmark for unveiling difficulties in social functioning, for example in neurodevelopmental condition such as ASD (Pavlova, 2012; Rosa Salva et al., 2015). Beyond the considerable success of this view (Kaiser & Pelphrey, 2012), refined theoretical models and robust quantitative approaches (i.e., meta-analysis) are lacking and whether and how BM processing is linked to social functioning is not clear. One possibility is to interpret BM processing as a building block of more complex social abilities in the context of the so-called “social brain”. Poor performance in BM discrimination in ASD would then be a marker of their social functioning difficulties (Kaiser & Pelphrey, 2012; Pavlova, 2012). The exact nature of the “social brain” construct (Elsabbagh & Johnson, 2016; Gammer et al., 2015; Whiten & van de Waal, 2017) and of other related constructs such as “theory of mind” or “mindreading” (Leslie, Friedman, & German, 2004; Meunier, 2017; Premack & Woodruff, 1978), which gained considerable attention in the last decades in a wide range of disciplines, remains however often vague and inconsistent among studies. The methodologies employed for exploring these constructs are often heterogeneous and related neurobiological underpinnings are unclear (Schaafsma, Pfaff, Spunt, & Adolphs, 2015).

## 2. Aim

As outlined in previous sections, BM perception is phylo- and onto-genetically robust and it has been hypothesized to be a putative marker for ASD (Di Giorgio et al., 2016; Klin et al., 2015; Pavlova, 2012). In addition, humans seem to have preserved a certain degree of confidence also in processing perturbed (or scrambled) versions of BM stimuli, both in temporal and spatial version (Troje & Westhoff, 2006). However, it is clear that different levels of information, such as global configuration and local dots trajectories, are involved in BM recognition.

Despite the fact that a rigorous selection of comparable studies for conducting a quantitative meta-analysis has the disadvantage of losing some information on the characterization of the BM phenomenon at large, this approach can help systematizing the heterogeneity of the findings reported so far. In our meta-analysis, we selected only studies that assessed the BM processing in ASD using PLD stimuli. As stated above, this kind of stimuli allows to investigate the essence of BM perception without the interference of other higher-level visual features that cannot be easily controlled for. We believe that focusing on low-level features is critical in our attempt to shed light on possible anomalies of BM processing in ASD. Moreover, this choice is consistent with other theoretical proposals which focused on findings from PLD stimuli to disentangle BM processing in the neurotypical population (Casile & Giese, 2005; Troje, 2008). To better understand BM processing anomalies in ASD, it is critical to investigate the impact of both the type of processing involved in different BM tasks and the type of stimuli that are used to assess this ability. These critical points will be addressed in the following subsections.

### 2.1 Deconstructing biological motion processing: a multi-level conceptual model

Given the high variability of the BM tasks employed with ASD individuals, we propose a conceptual classification of these tasks that aims at deconstructing the complex notion of BM in distinct (sub)components. The need for such deconstruction has been previously proposed also for other complex constructs such as “theory of mind” (Baron-Cohen, Campbell, Karmiloff-Smith, Grant, & Walker, 1995; Baron-Cohen, Leslie, & Frith, 1985; Fishman, Keown, Lincoln, Pineda, & Müller, 2014), “social brain” (Adolphs, 2009; Elsabbagh & Johnson, 2016) or “imitation” (Casartelli & Molteni, 2014; Casartelli & Parma, 2017; Caspers, Zilles, Laird, & Eickhoff, 2010; Lesourd et al., 2018). Indeed, when these constructs are treated as monolithic ones, we may run into misleading consequences. From a conceptual and epistemological perspective, they may convey different meanings and could translate into highly heterogeneous experimental protocols, which in turn can threat reproducibility of findings and reliability of results (Schaafsma et al., 2015).

In the literature, others have attempted to identify distinct (sub)components that play a role in BM processing. Troje (2008), for example, proposed a hierarchical model with four levels of processing with different complexity: 1) *life detector*, a cue of an animate object moving on the ground driven by the local motion, 2) structure-form-motion driven by the global motion, 3) action recognition and 4) style recognition, such as the recognition of emotional states. The author pointed out that the BM is a complex phenomenon characterized by a local spatio-temporal ballistic-velocity profile due to the gravity force, but that can also convey high-level information. Also, Casile and Giese (2005) proposed an interesting model that tries to explain the phylo- and ontogenetically highly preserved inclination in processing BM, showing the importance of the local motion in BM recognition. The authors hypothesized a hierarchy of neural detectors that extract motion features with different complexity of BM: 1) local motion energy detectors, 2) detectors for horizontal and vertical opponent motion, 3) detectors for complex global optic flow patterns and 4) detectors for complete biological motion patterns. Both these models highlighted a clear hierarchical organization in BM processing. Although these models have the merit of focusing on the need of deconstructing BM first, for our specific aim of better understanding BM anomalies in ASD, we propose here a new distinction between different levels of BM processing according to task complexity. We hypothesize that these distinct levels of BM processing could be differently affected in participants with ASD. Specifically, we propose a three-level model that distinguishes between: I) first-order processing of BM, II) direct processing of BM, and III) instrumental processing of BM.

> **I) First-order processing of BM** refers to the implicit detection of BM that implicates a behavioral response which does not entail any explicit recognition or categorization of BM. Experimenters infer that participants have detected a BM stimulus by simply evaluating their spontaneous behavioral response (e.g., looking time in preferential looking paradigms).
>
> **II) Direct processing of BM** involves an explicit discrimination between distinct features of the stimuli, using tasks with different types of BM stimuli (e.g. Rightward Vs. Leftward walker), or an explicit detection and/or recognition of BM that employs BM stimuli and non-BM stimuli (e.g., Human Vs. Vehicle; Biological Vs. its Scrambled version).
>
> **III) Instrumental processing of BM** implicates that the observer takes advantage of the processing of a BM stimulus for a secondary purpose, i.e. to disclose distinct and potentially more complex information such as inferring about other’s intentionality, emotional states or actions that are conveyed in the BM stimulus.

A meta-analysis based on this three-level model allows to disentangle whether different levels of processing of the rich information conveyed by PLD-based BM could be at least partially the source of the variability of the findings of BM processing reported in ASD.

### 2.2 Do different features of BM stimuli imply different performance?

Another possible source of variability in the BM processing results in the ASD population may stem from the fact that BM is characterized by a moving structure that requires both spatial and temporal binding of information. Interestingly, Casile and Giese (2005) found that local motion seems to be the most important feature leading to robust BM processing. Indeed, they showed that even if the spatial arrangement of the PLD dots is inconsistent with the human skeleton, a plausible local optic flow is enough to suggest the presence of a BM stimulus.

Given the evidence of anomalies in low-level features processing in individuals with ASD (Robertson & Baron-Cohen, 2017), we think that it could be extremely useful to test if the data heterogeneity showed by ASD individuals in BM processing can be driven by the manipulation of different stimulus features. In other words, perceptual anomalies that are typically associated with ASD could potentially impact on BM processing more strongly when certain type of stimuli are employed. Therefore, the present work aims to preliminarily assess whether the difference between the performance of ASD and TD individuals is modulated by the type of low-level manipulation (spatial vs. temporal) of scrambled PLD stimuli used as comparison. Moreover, to assess whether spatio-temporal processing anomalies are specific to BM stimuli – in virtue of their peculiar pattern – or they are generalized, we tested if there is a difference between the performance of ASD and TD individuals between BM and non-BM stimuli.

Altogether, the present meta-analysis will compare the performance of ASD and TD groups according to the following four steps:

1. A general analysis exploring all studies using PLD to investigate BM in ASD (***Biological motion in ASD vs. TD*** [***1=All BM studies***]);
2. An analysis testing BM processing in ASD according to our three-levels model (i.e., first-order, direct and instrumental BM processing) (***Level of processing of biological motion in ASD vs. TD*** [***2=Level of processing***]);
3. An analysis investigating the contribution of spatial and temporal features in how individuals with ASD process BM (***Impact of low-level features of BM scrambled stimuli in ASD vs. TD*** [***3=Low-level features***]);
4. An analysis testing ASD performance in non-BM stimuli (***Non-biological motion in ASD vs. TD*** [***4=Non-BM***]).

## 3. Methods and Results

### 3.1 Literature Search

Following the PRISMA-P guidelines (Moher et al., 2015), we have established a protocol detailing the a priori rationale, the methodological and analytical approaches to which we adhered during the preparation of the present meta-analysis. This protocol does not represent an update of any other reports and it is not part of a systematic review on this topic.

An automatic literature search began on September 2017 and the last search was performed in March 2018. We searched for all studies published in English in both PubMed and PsycInfo, by performing a search on titles, abstracts and keywords. For direct replication, we include the three Boolean queries used to the search the PubMed database: 1) ((“biology”[MeSH Terms] OR “biology”[All Fields] OR “biological”[All Fields]) AND (“motion”[MeSH Terms] OR “motion”[All Fields])) AND (“autistic disorder”[MeSH Terms] OR (“autistic”[All Fields] AND “disorder”[All Fields]) OR “autistic disorder”[All Fields] OR “autism”[All Fields]); 2) “Human motion”[All Fields] AND (“autistic disorder”[MeSH Terms] OR (“autistic”[All Fields] AND “disorder”[All Fields]) OR “autistic disorder”[All Fields] OR “autism”[All Fields]); 3) point-light[All Fields] AND (“autistic disorder”[MeSH Terms] OR (“autistic”[All Fields] AND “disorder”[All Fields]) OR “autistic disorder”[All Fields] OR “autism”[All Fields]). In addition, we also performed a manual literature search which encompassed a search of the reference lists of the review articles and the original research papers retrieved via the Boolean queries. One paper was retrieved by using this method. Altogether the automatic and the manual literature searches resulted in 114 unduplicated published articles. No reviews, book chapters, master and doctoral theses, conference presentations or unpublished data were hereby included.

### 3.2 Screening Process

The studies retrieved from the automatic and manual literature searches were independently assessed and screened by two authors (A.F. and L.Ra.). At the end of the first round of screening, they evaluated together the differences emerged from the two separate searches and discussed with the other authors the final set of papers to be included, based on the inclusion criteria as detailed below. The *n* reported represents the number of studies excluded at each step. Please, also refer to Figure 2.
1. Screening: from the analysis of title and abstract, the empirical studies that did not include a task in which the processing of BM was assessed via PLD in both an ASD and a TD group were excluded from the initial paper count (*n*=55).
2. Eligibility: from reading the whole publication, studies were further excluded if they did not meet the same criteria of the previous screening step (*n*=16).
3. Qualitative assessment: from reading the whole publication, the articles were further excluded if they did not meet the following criteria:

a. If the study sample was the subset of a bigger sample presented in a different paper, the study with the smaller sample was discarded (*n*=2; i.e. Klin & Jones, 2008; Koldewyn, Whitney, & Rivera, 2011).
b. If the study did not investigate behavioral measures such as accuracy, sensitivity (d’), threshold estimation, reaction times (RTs), or percentage of preferential looking (or first orienting) in a BM task. Thus, all the studies that use a passive view task in order to investigate BM perception using EEG or fMRI were discarded (*n*=8).
c. If the dependent variables and/or task employed were not comparable to the majority of the studies included. Specifically, these three studies were removed for the following reasons: i) Krüger and colleagues (2017) used a confidence rating as the dependent variable, no accuracy or RTs were reported; ii) Van Boxtel and colleagues (2016) used as the dependent variable the difference in the point of subjective equality (PSE) between the walking and running conditions, and thus the data related to BM performance could not be retrieved; iii) Swettenham and colleagues (2013) used an attentional cuing paradigm in which cues were BM stimuli and reported RTs for valid and invalid conditions, which do not primarily map onto BM processes.
d. If the study did not have sufficient statistical information (either present in the paper, supplementary materials or provided after contacting the authors) (*n*=3).

**Figure 2.**
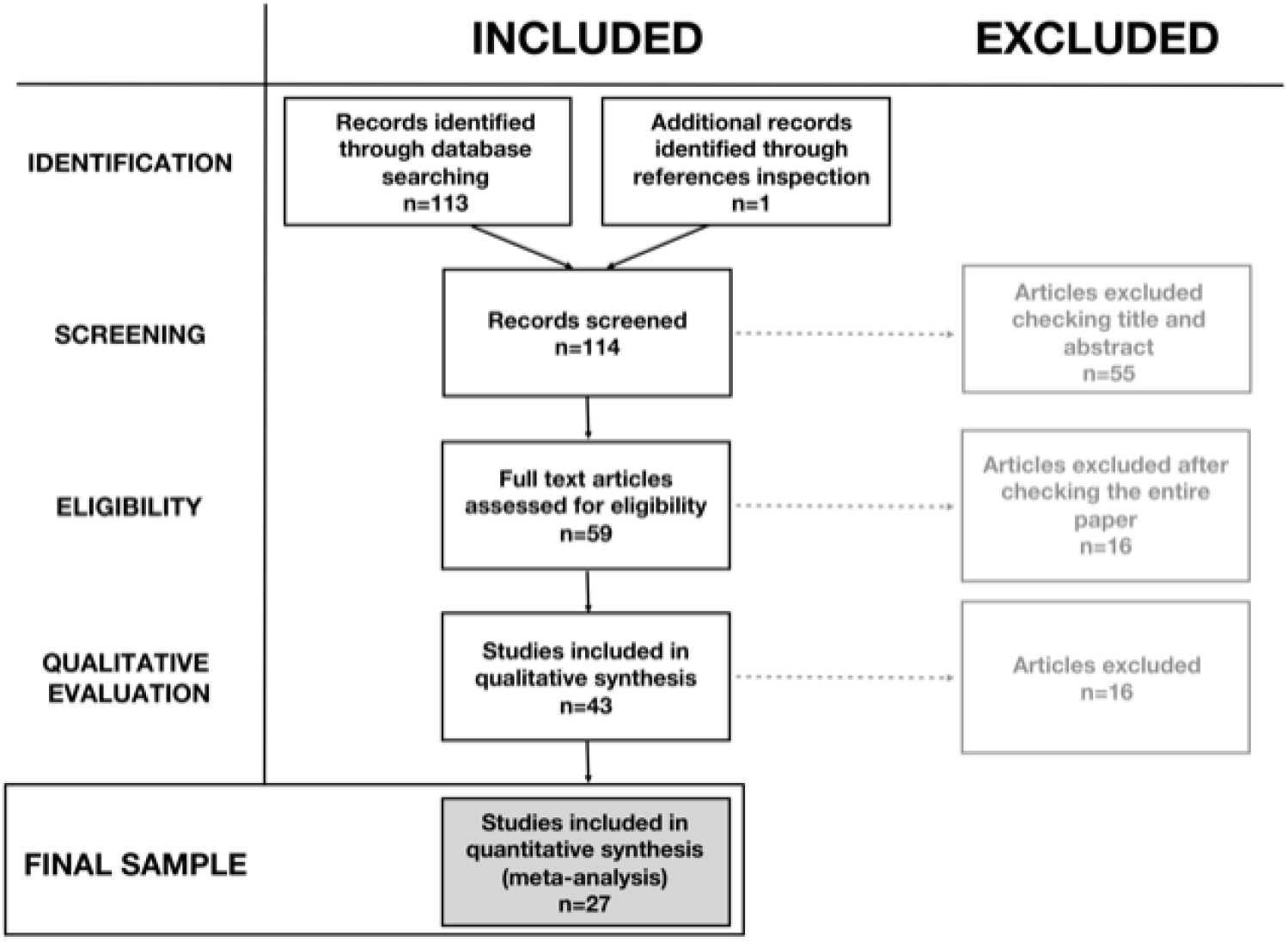
Flow diagram of the selection of studies included in the current meta-analysis. The number of studies included or excluded at each step of the evaluation process is indicated in each of the diagram boxes.

### 3.3 Coding and Reliability

The final sample of studies were coded based on the following rules:

1. Level of BM processing, which led to the creation of three main categories; 1) First-order; 2) Direct recognition; 3) Instrumental recognition of BM;
2. Type of scrambled stimulus used in tasks that assess the perception of BM vs. its scrambled version: 1) no scrambled; 2) spatial; 3) temporal; 4) spatial and temporal scrambled.
3. Type of motion stimuli presented: 1) BM, when the data refer to the performance following the presentation of solely BM stimuli; 2) BM vs. non-BM, when the dependent variable refers to a task in which the perception of BM stimuli is contrasted with the perception of non-BM stimuli; 3) non-BM, when it is possible to separately extract data concerning only the performance of non-BM stimuli (i.e., the scrambled version of the BM stimulus or the motion of an object, such as a truck or a bicycle). Disagreement and inconsistencies between the two raters (A.F. and L.Ra., N=7/114, 6% of all assessments) were questioned with all authors to gather consensus.

In addition, for all the studies we collected:

1. The relevant sample sizes of both ASD and the TD groups;
2. The percentage of male participants in each of the two groups;
3. The mean age of each of the two groups;
4. The test employed to assess general cognitive level of participants, and the relative mean score for each of the two groups;
5. The type of behavioral measure collected: accuracy rates (including error rates) or percentage of looking time. Since RTs were present only in 3 of the studies included in this meta-analysis, we preferred to exclude them to increase the homogeneity and the comparability of measures;
6. Whether behavioral tasks are performed during an EEG or fMRI acquisition;
7. A brief qualitative description of the task and the displayed stimuli.

All this relevant information about the studies selected to be included in the final analyses is reported in Table 1. Moreover, to calculate the Cohen’s *ds*, descriptive statistics of the ASD and TD groups’ performances or the associated test statistics were extracted. If further groups were tested in the study, then the statistics relative to the other groups were not considered.

**Table 1.**
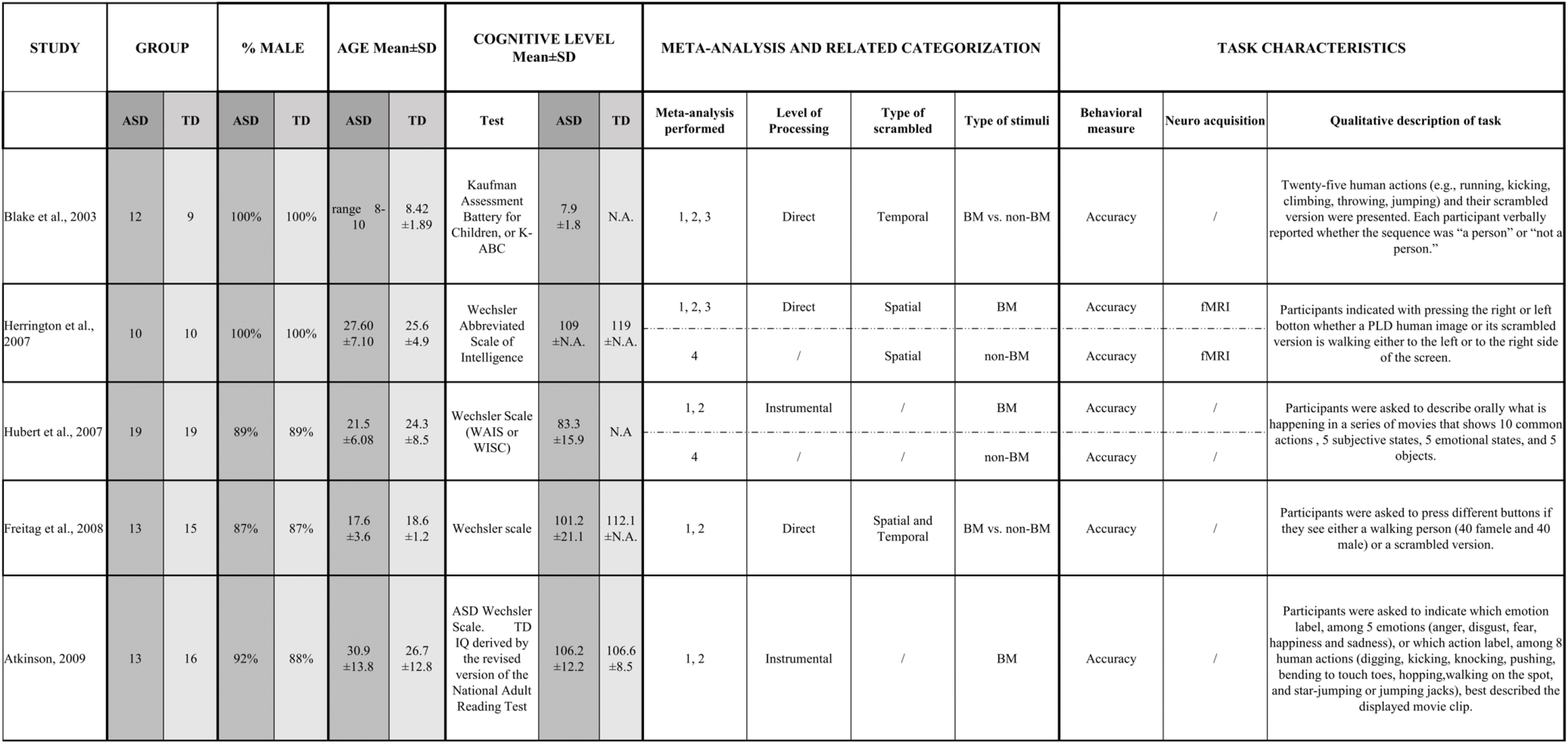
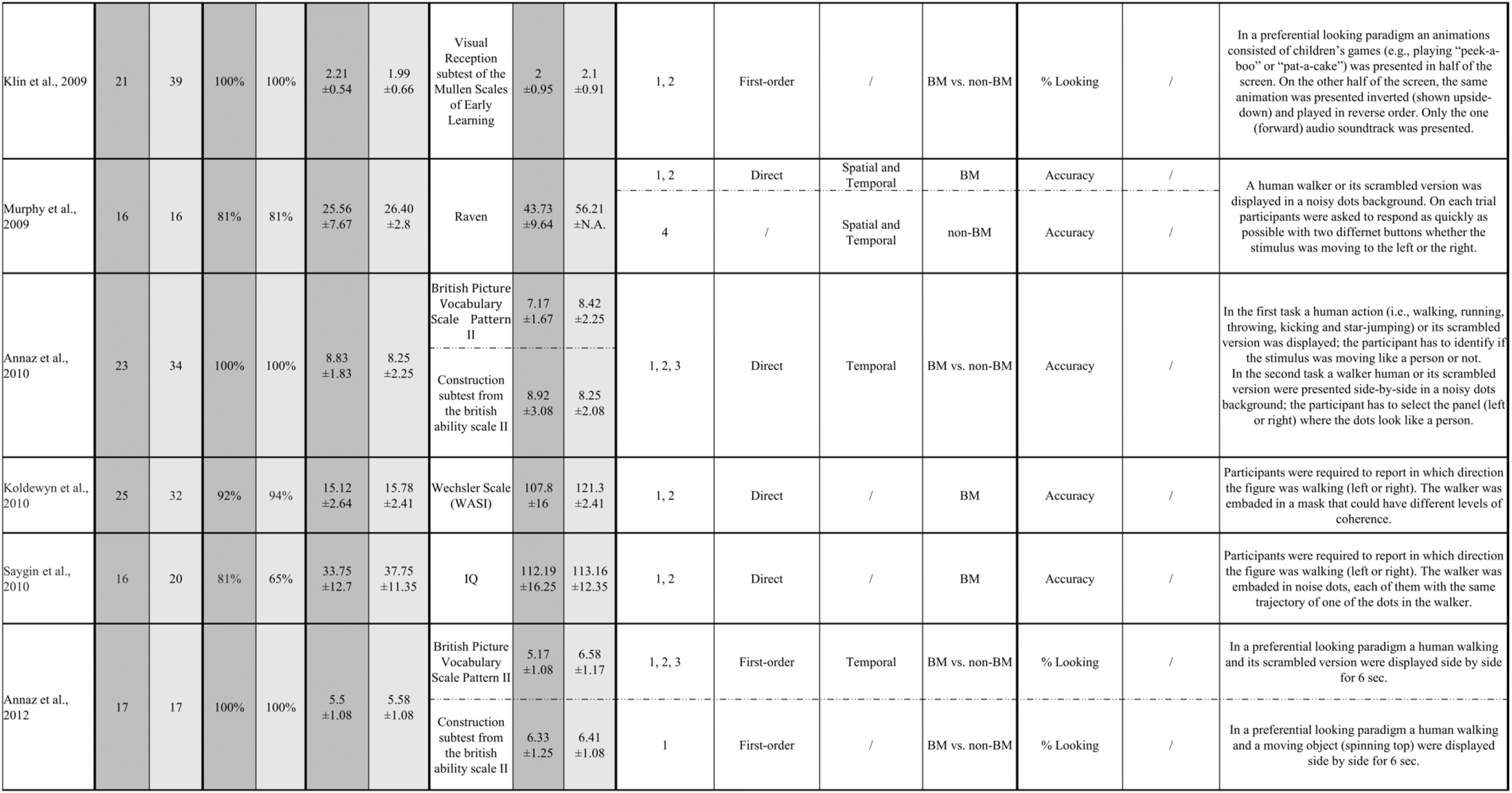
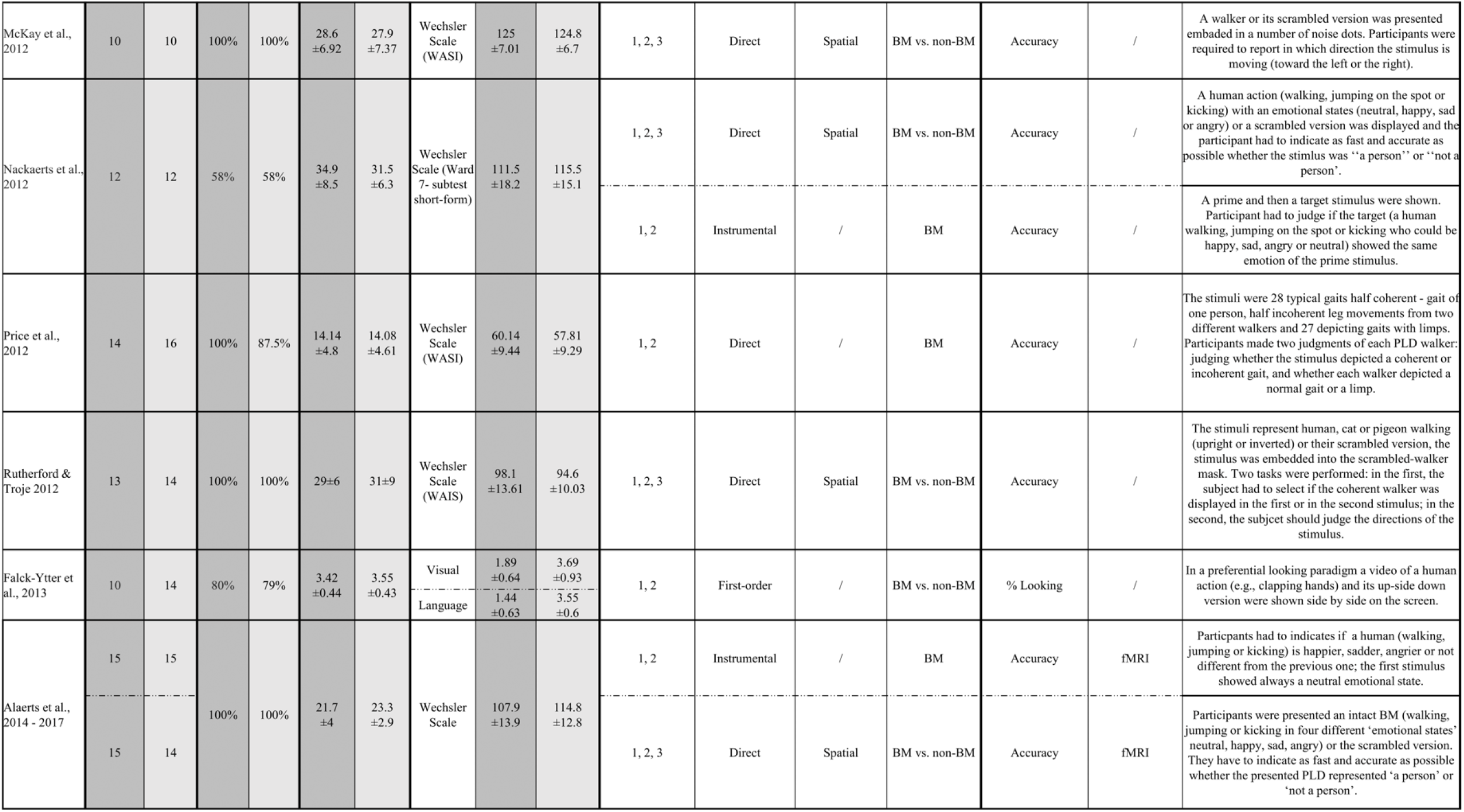
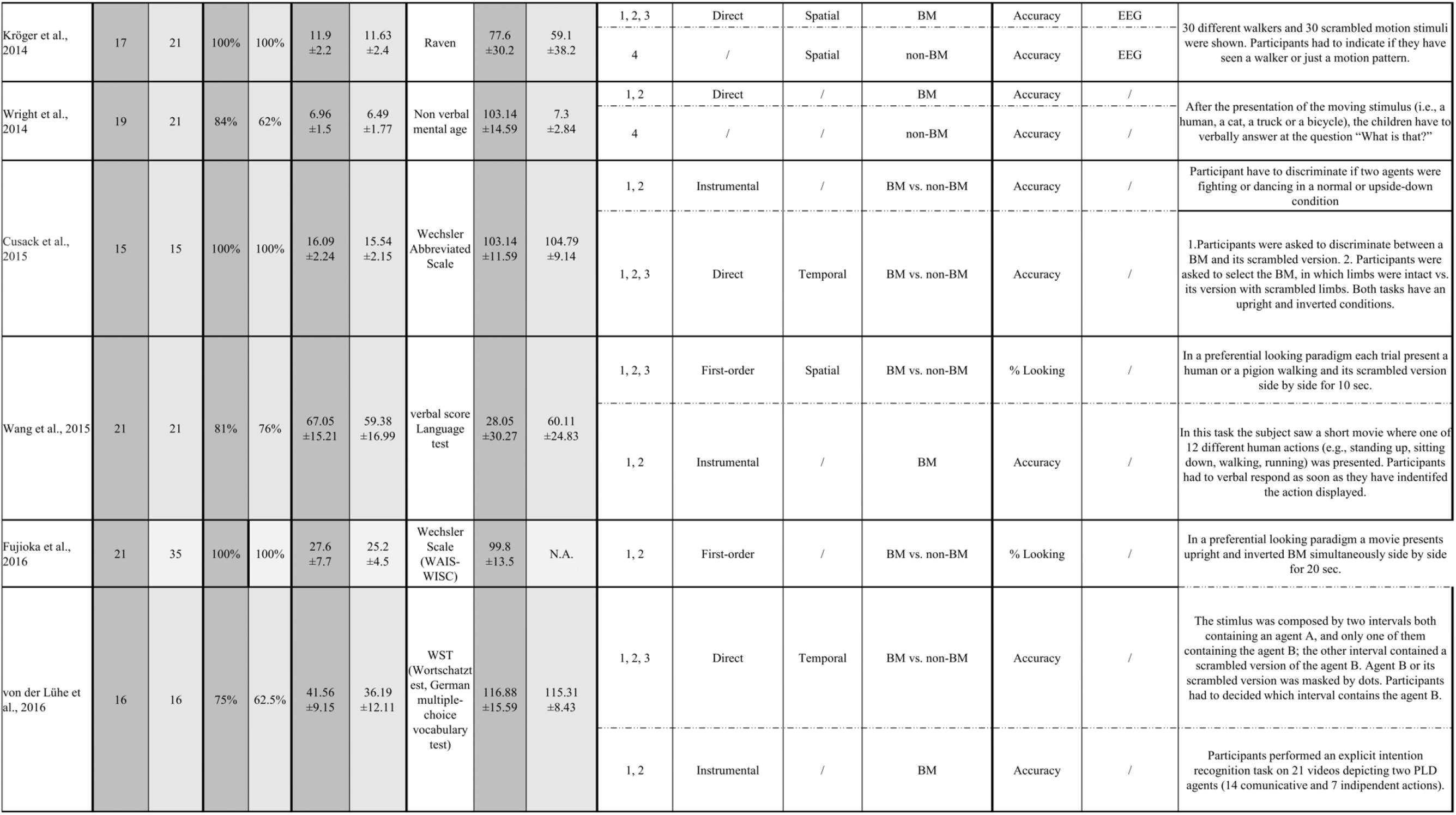
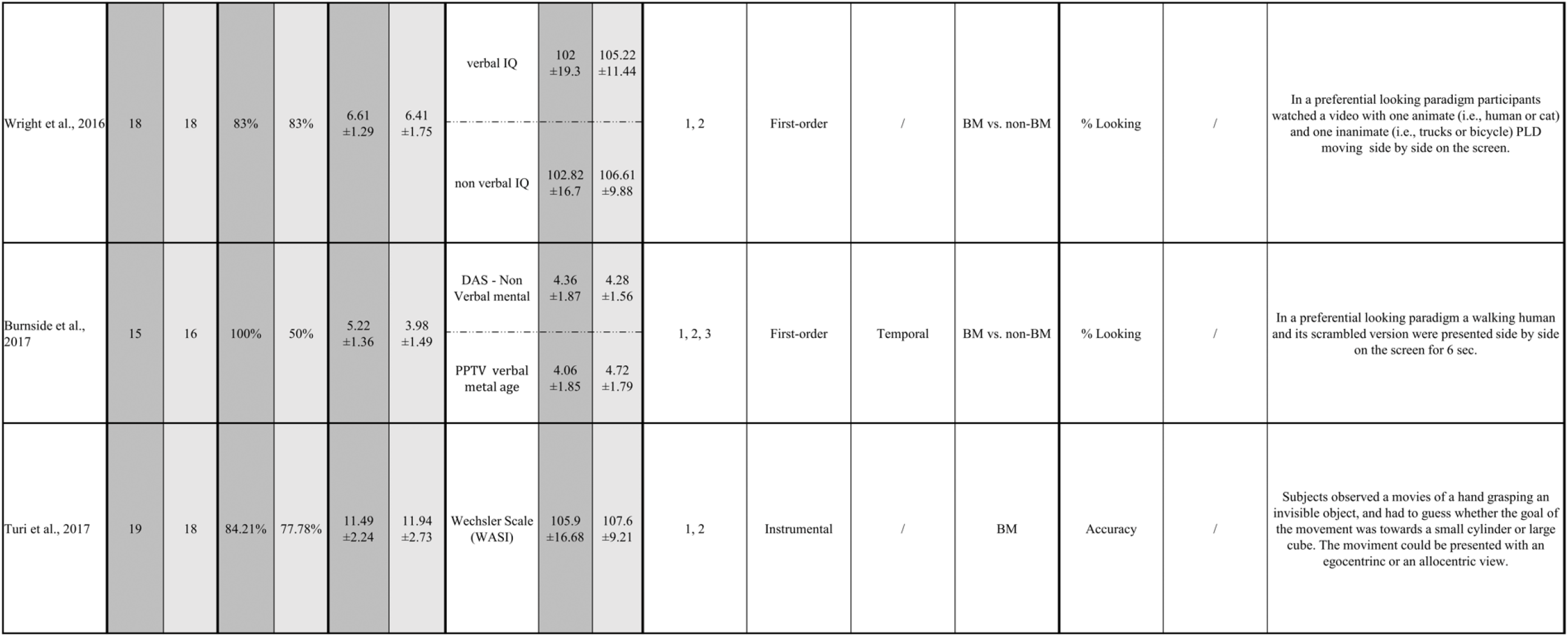
The table contains the final set of studies included in the meta-analysis. For each study descriptive statistics are reported for both ASD and TD groups (since different tests were used in order to assess the participants’ cognitive level, in the subcolumn ‘Test’ the name of the test is specified). When the data were not availaible in the paper N.A.is reported. The column ‘Meta-analysis and related categoization’ contains four subcolumns where we reported in which meta-analysis each study was included (1=All BM studies, 2=Level of processing, 3=Low-level features, 4=Non-BM), and the relative categorization: Level of processing(First-order, Direct, Instrumental), Type of scrambled (Spatial, Temporal, Spatial and Temporal) and Type of stimuli(BM=only biological motion stimuli, non-BM=only non-Biological motion stimuli, BM vs. non-BM when a biological stimulus has to be discriminated from a non-biological one). In the ‘Task characteristics’ column we reported which behavioral measure was assessed in the task (Accuracy, % Looking), if the task was performed during an EEG or a fMRI acquisition, and a brief description of the implemented task.

### 3.4 Meta-Analytic Procedures

#### 3.4.1 Effect size calculation

For each observation, using the descriptive or test statistics present in the papers selected, we calculated Cohen’s *d* as a measure of the size of the group effect (ASD vs. TD) on the variable of interest. Only two studies explicitly reported Cohen’s *d*. For all other studies, we extracted Cohen’s *d* from mean and standard deviations (or standard errors), or from the *t*-test value (or the associated *p*-value) of the reported statistical analyses. The mean and the standard deviation were available in most of the studies, therefore we used the following equation (1) to compute the Cohen’s *d*^1^ (see (Borenstein, Hedges, Higgins, & Rothstein, 2009; Lakens, 2013):

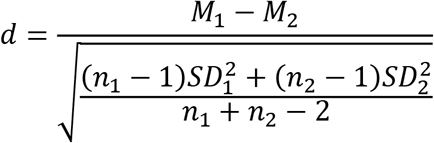

where *M*_1_ is the mean of the dependent variable for the first group (e.g., ASD), *M*_2_ is the mean of the dependent variable for the second group (e.g., TD), *SD*_1_ and *SD*_2_ are respectively the standard deviations of the mean for the first and the second group, *n*_1_ and *n*_2_ refer to the number of participants in the first and the second group, respectively. We arbitrarily set the direction as positive indicating a better performance for the TD group as compared to the performance of the ASD group (e.g., lower error rate). According to the guidelines of Cohen (1988), an absolute effect size of 0.2-0.3 is considered a small effect, ∼0.5 a medium effect and >0.8 a large effect. The variance associated to each Cohen’s *d* was computed following equation (2) (Borenstein et al., 2009):

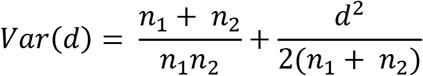

In some cases, two or more Cohen’s *d*s referring to the same study were included in the same meta-analysis (see below). When more than one task was included in the same (sub)group of the meta-analysis, we pooled those Cohen’s *d*s in order to compensate for the possible underestimation of the variance of the mean effect size resulting from the inclusion of multiple effect sizes from a single study in the same meta-analysis (see Borenstein et al., 2009; Tanner-Smith, Tipton, & Polanin, 2016). The Cohen’s *d*s referring to the same study are pooled using the formulas reported by Borenstein et al., 2009, p. 230 (equations 3 and 4):

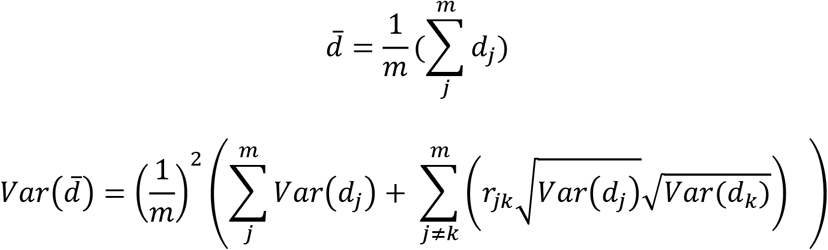

where 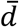 and 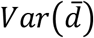 are the average Cohen’s *d* and its variance, *m* is the number of Cohen’s *d*s that were pooled, *d*_*j*_and *d*_*k*_are the Cohen’s *d*s referring to measures *j* and *k*, and *r*_*jk*_ is the correlation between measures *j* and *k*, which was arbitrarily set to 0.5^2^. The maximum number of Cohen’s *d*s that were pooled is 10. This number varies depending on the meta-analysis performed.

In each of the four meta-analyses (see below), multiple Cohen’s *d*s from the same study were pooled when they referred to tasks that were coded in the same manner. For instance, in (Annaz et al., 2010) there are two tasks, one measuring the sensitivity (d’) for the perception of BM in normal PLD vs. scrambled stimuli, and the other establishing the thresholds for the detection of a PLD walker in noise. Since both tasks were coded as *direct processing*, the two Cohen’s *d* were always pooled together. This was also the case of the “action”, “subjective state”, and “emotion” conditions in Hubert et al. (2007), that were all coded as tasks of *instrumental processing*. In some cases, multiple Cohen’s *d*s from the same study could be kept separate or pooled together depending on whether the categorical distinction between the tasks was relevant or irrelevant for the meta-analysis at hand. For instance, in Nackaerts et al. (2012) the Cohen’s *d*s referring to the task categorized as *direct processing* and the task categorized as *instrumental processing* were pooled in the first general meta-analysis [*1=All BM studies*], but they were kept separate in the meta-analysis testing the effects of the level of BM *processing* [*2=Level of processing*].

#### 3.4.2 Mixed effects models

Statistical analyses were conducted in R, using the package *metafor* (Viechtbauer & Cheung, 2010). The meta-analysis performed was analyzed using a mixed effects model (Viechtbauer & Cheung, 2010), estimating the regression coefficients for each moderator included as well as the error between experiments. No within error estimate was included in the model, because we only included one Cohen’s *d* per study. The rationale for this choice was that the majority of data from the studies presented a single effect size and a traditional random-effect model would suffice (Borenstein et al., 2009; see also Note 2). A simple model with one moderator variable is given in equation (5):

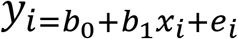

for *i* ∈ {1,…,*n*}

*y_i_* is the Cohen’s *d* for the *i*^th^ study

*b_0_* is the fixed intercept for the regression model

*b_1_* is the fixed slope for the regression model

*x_i_* is the predictor for the *i*^th^ study

*e_i_* ∼ N(0, σ) is a Gaussian error term.

The variance components of the model including the predictor variables refer to the unexplained variance. To answer specific questions, we run four separate meta-analyses:

1. ***Biological motion in ASD vs. TD*** [***1=All BM studies***]. We included all the studies that were selected, and, for each of them, a single Cohen’s *d* was inserted (n=26). Since the cognitive functioning was assessed with different tests across studies, as reported in Table 1, we computed a Cohen’s *d* per study to have a comparable measure of the cognitive level to be used as a possible moderator. A positive Cohen’s *d* indicates that the cognitive level of the TD group is greater than the cognitive level of the ASD group. We included this moderator in the meta-analytic models as a way to assess if the ASD diagnosis has a specific effect on the processing of BM, which cannot simply be attributed to a reduction in cognitive level. Moreover, we insert also the mean age of ASD and of TD groups as moderators in order to control for the possible effect of age.
2. ***Level of processing of biological motion in ASD vs. TD [2=Level of processing].*** We included one Cohen’s *d* per study in each of the three levels of BM processing identified (n=7 first-order, n=16 direct, n=8 instrumental). ‘Level of processing’ was included in the meta-regression analysis as a categorical predictor, and no further moderating effects were considered here, due to the limited number of studies included in each group.
3. ***Impact of low-level features of BM scrambled stimuli in ASD vs. TD*** [***3=Low-level features***]. We included one Cohen’s *d* for each study that used the scrambled version of the BM stimulus (n=7 spatial, n=6 temporal). We considered two types of scrambled stimuli according to the manipulation, spatial or temporal, employed to generate them. To this aim, this meta-analysis includes a categorical moderator, ‘type of scrambled’, which consists of two levels: 1) *spatial scrambled*, the difference between the scrambled and the BM stimulus is limited to a difference in their spatial features. In other words, in the scrambled version each dot of the PLD was randomly displaced in a different spatial position; 2) *temporal scrambled*, the difference between the scrambled and the BM stimulus is limited to a difference in their temporal features. In other words, each dot in the PLD scrambled stimulus was presented in the same spatial position but the motion of each single dot was temporally out of phase as compared to the BM stimulus. Studies with both spatial and temporal manipulations were excluded since they were not numerous enough to create a stand-alone group (n=2). Considering the limited number of studies per level, we refrained from including other moderators.
4. ***Non-biological motion in ASD vs. TD*** [***4=Non-BM***]. This meta-analysis is limited to the studies that test, together with the perception of a BM stimulus, also the processing of another stimulus that can be defined as a non-BM stimulus (i.e., an object as a bike or a truck presented in PLD; (Hubert et al., 2007; Wright, Kelley, & Poulin-Dubois, 2014) or of a scrambled version of the BM stimulus (Herrington et al., 2007; Kröger et al., 2014; Murphy, Brady, Fitzgerald, & Troje, 2009) (n=5). It is important to note that we do not consider this analysis to be ultimately conclusive, because an extensive analysis of this type should include also other types of non-BM tasks such as coherent motion, form-from-motion, etc. However, we still believe that this aspect is worth of investigation to provide an initial characterization of possible dissociation between BM and non-BM processing in ASD vs. TD.

All tests were conducted with a significance level of 5%. The results will be presented including the following measures: weighted mean effect sizes (Cohen’s *d*), 95% confidence intervals, *I*^2^ heterogeneity values, and *p* values. All error bars in forest plots are 95% confidence intervals and were generated by applying customized R scripts.

#### 3.4.3 Moderator correlations

Being aware of the need to control for confounded moderators (Field & Gillett, 2010), we opted to use ANOVAs (performed in R with the *aov* function) in order to evaluate possible systematic relationships between the categorical predictor (i.e. ‘level of processing’, ‘type of scrambled’) and the quantitative predictor (i.e. ‘cognitive level’, ‘mean age of ASD’, ‘mean age of TD’). This procedure allowed to define whether the cognitive level or the groups age differed systematically across the *levels of processing* and *type of scrambled*.

#### 3.4.4 Publication bias

To determine the presence of publication bias for the main effect of each meta-analysis, we plot each study’s effect size (Cohen’s *d* ASD vs. TD) against its own standard error in a funnel plot. Publication bias is computed as a negative association between the sample size and the effect size across studies, since to be significant a result coming from a small sample must have a large effect size. More specifically, publication bias occurs when studies with small samples are more likely to be published when they report statistically significant results (i.e., a large effect size), as compared to when they do not (i.e., a small effect size). The publication bias implies an overestimation of the true effect size, for which appropriate correction methods are available (see below).

### 3.5 Meta-analysis results

#### 3.5.1 Biological motion in ASD vs. TD [1=All BM studies]

The analysis including all papers contrasting the processing of biological motion in ASD vs. TD revealed a medium effect of the difference in performance between the two groups, Cohen’s *d* = 0.57, *SE* = 0.14, CI 95% = [0.30, 0.85], *z* = 4.08, *p* <.0001. In other words, there is a medium effect size indicating that individuals with ASD have more difficulties in BM processing than TD individuals (Figure 3). However, the *I*² coefficient reveals that 79% of the variability of the effect sizes can be attributed to the heterogeneity among the true effects (Test for Heterogeneity: *Q*_(25)_ = 108.89, *p* <.0001). Cognitive level, mean age of ASD group and mean age of TD group proved to have no statistically significant effect on Cohen’s *d* (cognitive level: *b* = 0.23, *SE* = 0.24, CI 95% = [-0.23, 0.70], *z* = 0.98, *p* =.33; mean age of ASD group: *b* = 0.003, *SE* = 0.01, CI 95% = [-0.02, 0.02], *z* = 0.3, *p* =.76; mean age of TD group: *b*= 0.002, *SE*= 0.01, CI 95%= [-0.02, 0.02], *z* = 0.14, *p* =.89).

**Figure 3.**
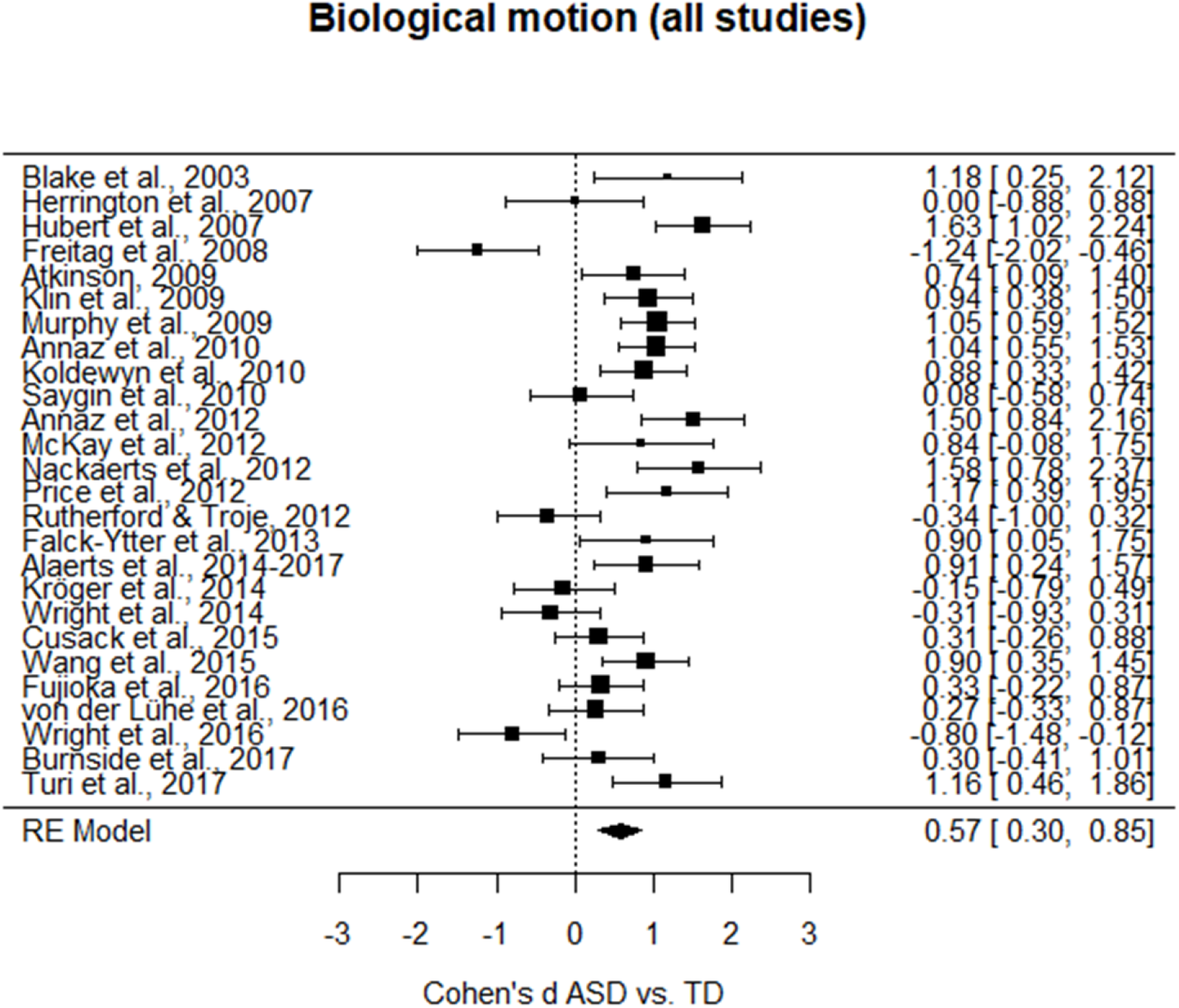
Forest plot of the differences in the BM performance between the ASD and TD groups. Positive values indicate a better performance for the TD group as compared to the performance of the ASD group.

A visual inspection of the funnel plot resulting from this meta-analysis suggests a quite symmetrical distribution of the studies. To confirm whether or not a publication bias is present in this case, we computed a trim and fill test (Taylor & Tweedie, 1998). Such test estimates the absence of three studies on the left side (*SE* = 3.39) and none on the right side. When computing a new average effect size including the effect sizes of the three studies estimated to be absent, the average effect size was still significantly different from zero, Cohen’s *d* = 0.45, *SE* = 0.15, CI 95% = [0.16, 0.73], *z* = 3.06, *p* =.002 (Figure 4).

**Figure 4.**
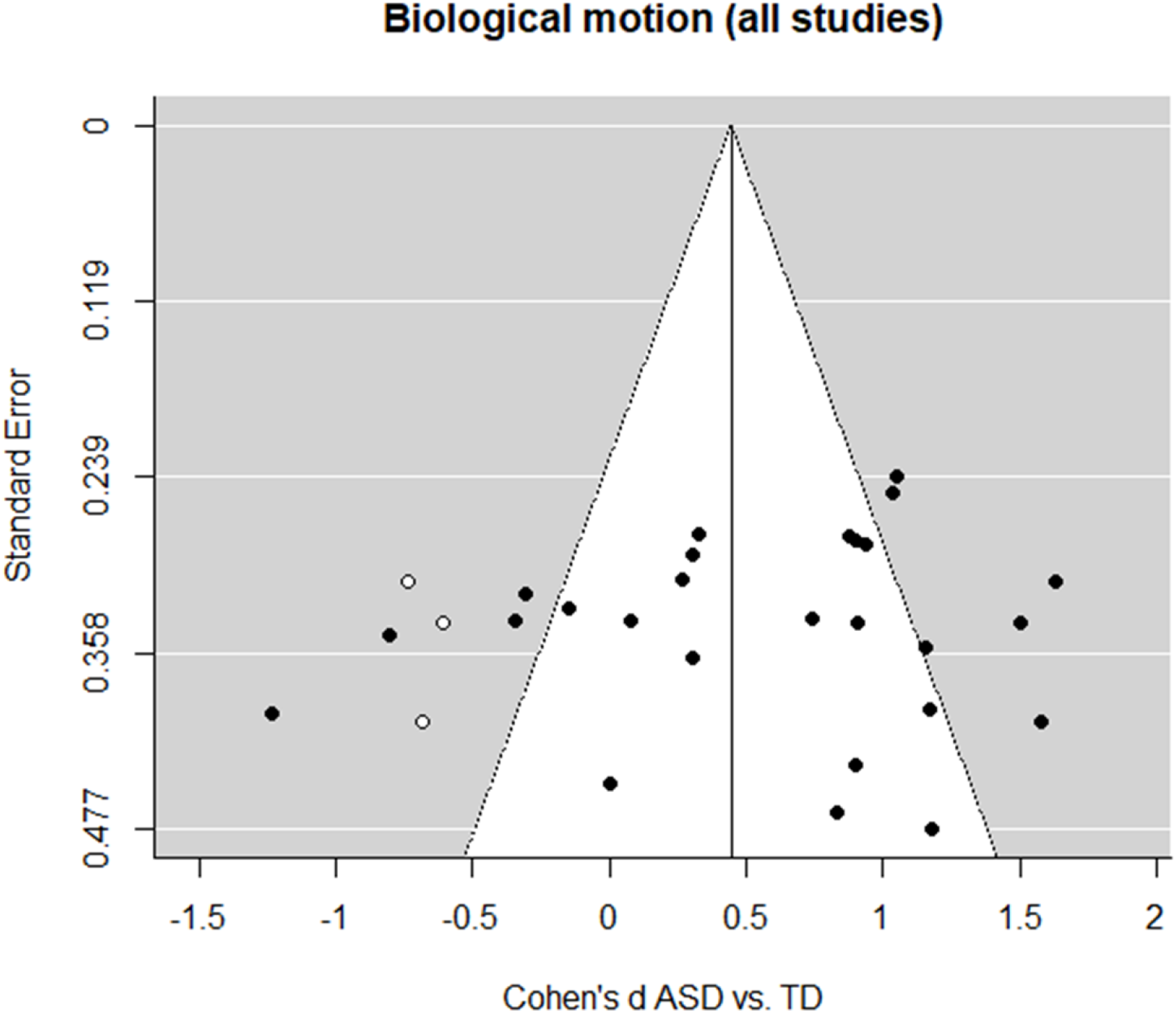
Funnel plot indicating the publication bias in studies of BM processing in ASD vs. TD. White circles represent the missing studies identified by the trim and fill test.

Thus, since the effect size was still medium and significant even after the inclusion of the estimated missed studies t, the difference in the processing of BM between ASD and TD groups cannot be considered dependent on publication bias.

#### 3.5.2 Level of processing of biological motion in ASD vs. TD [2=Level of processing]

To investigate the role of the level of processing needed to accurately perceive BM in ASD vs. TD, we computed a mixed effect model with ‘level of processing’ as a moderator. This moderator has three levels: i) first-order processing; ii) direct processing; iii) instrumental processing, indicating the three subgroups in which tasks were divided. The *I*^2^ coefficient indicates that the heterogeneity among the true effects is ∼73% (Test for Residual Heterogeneity: *QE*_(28)_ = 103.03, *p* <.0001). The results reveal a significant effect of the moderator, *QM*_(3)_ = 29.61, *p* <.0001. Thus, to further explore each level of the moderator we calculated separate meta-analyses for each level of processing. When analyzing the *first-order processing* level, we obtained Cohen’s *d* = 0.55, *SE* = 0.27, CI 95% = [0.02, 1.09], *z* = 2.04, *p* =.04, and the heterogeneity was comparable to that of the full model (*I*^2^ = 79%, *Q*_(6)_ = 27.04, *p* <.0001, Figure 5). When analyzing the *direct processing* level, we obtained Cohen’s *d* = 0.40, *SE* = 0.18, CI 95% = [0.05, 0.74], *z* = 2.24, *p* =.03, and the heterogeneity was comparable to that of the full model (*I*^2^ = 76%, *Q*_(15)_ = 61.29, *p* <.0001, Figure 6). Instead, the *instrumental processing* level is the one showing the greatest difference in the processing of BM between ASD and TD, Cohen’s *d* = 1.04, *SE* = 0.19, CI 95% = [0.68, 1.40], *z* = 5.62, *p* <.0001, and the heterogeneity was reduced as compared to that of the full model (*I*^2^ = 52%, *Q*_(7)_ = 14.70, *p* =.04, Figure 7).

**Figure 5.**
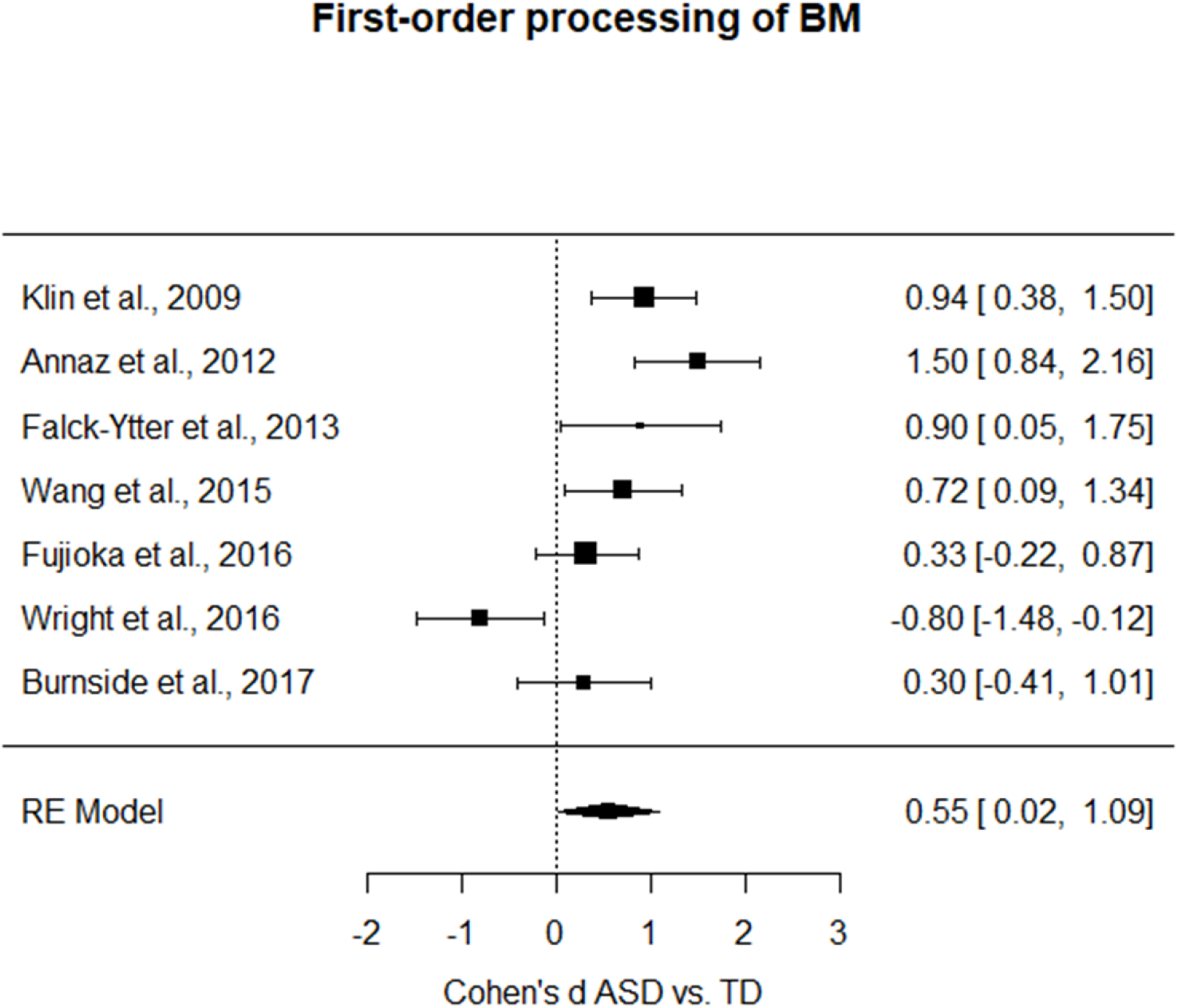
Forest plot of studies investigating the difference in first-order level of BM processing between ASD and TD group (i.e. positive values indicate a better performance for the TD group as compared to the performance of the ASD group).

**Figure 6.**
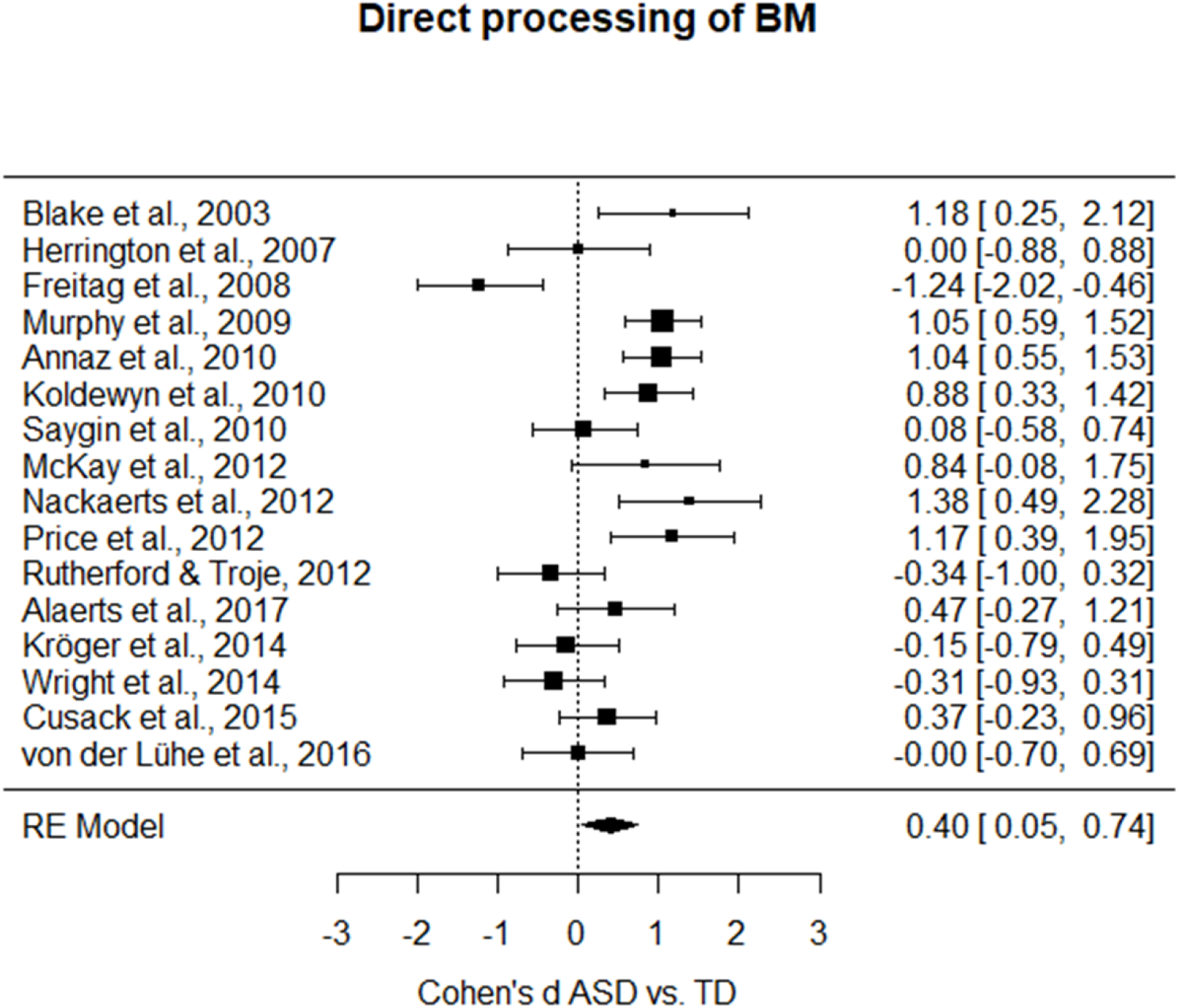
Forest plot of studies investigating the difference in direct level of BM processing between ASD and TD group (i.e. positive values indicate a better performance for the TD group as compared to the performance of the ASD group).

**Figure 7.**
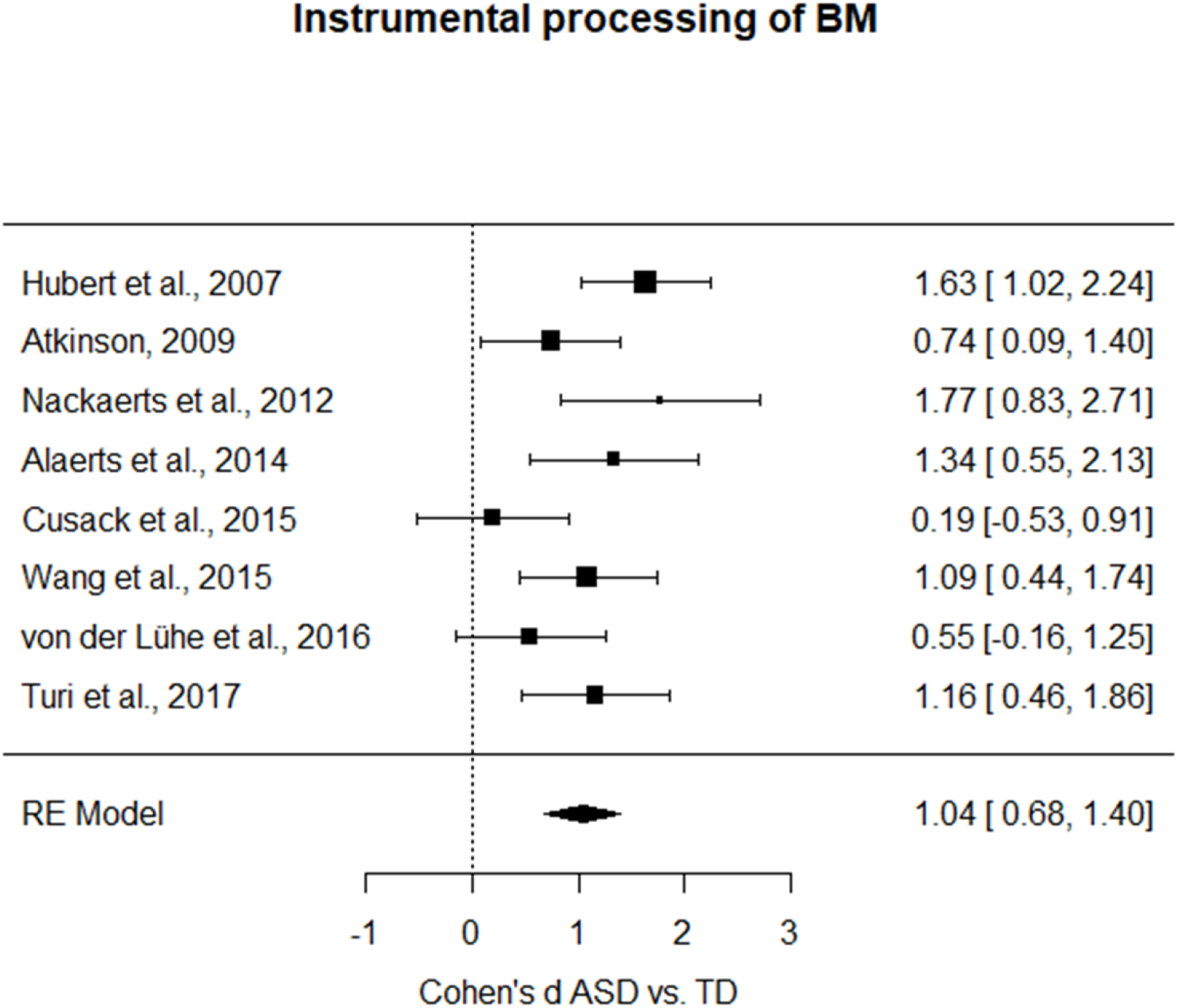
Forest plot of studies investigating the difference in instrumental level of BM processing between ASD and TD group (i.e. positive values indicate a better performance for the TD group as compared to the performance of the ASD group).

Moreover, in order to test if there is a significant difference between the magnitude of these estimated effect sizes, we performed three *z*-tests between the Cohen’s *d*s of the different levels of processing using the Bonferroni correction method. Only the difference between *direct* and *instrumental* level was statistically significant (direct vs. instrumental: *z* = −2.53, *p* =.03; first-order vs direct: *z*= −0.46, *p* ∼ 1; first-order vs. instrumental: *z* = −1.48, *p* =.42). To evaluate possible systematic differences in cognitive level across the levels of BM processing, we performed a one-way ANOVA on the Cohen’s *d*s referring to the cognitive level with ‘level of processing’ as a factor. No statistically significant differences were retrieved (*F*_(1, 24)_ = 2.11, *p* = .16, *η*_p_ ^2^ = 0.08). Moreover, two one-way ANOVAs showed that neither the mean age of the ASD groups nor that of the TD groups differed systematically across the levels of processing (*F*_(1, 28)_ = 2.87, *p* =.10, *η_p_*^2^ = 0.09, and *F*_(1, 29)_ = 3.45, *p* =.07, *η_p_*^2^ = 0.1, respectively). In sum, the effects of level of processing on Cohen’s *d* cannot be explained in terms of systematic differences between cognitive level, mean age of the ASD groups, or mean age of the TD groups.

#### 3.5.3 Impact of low-level features of BM scrambled stimuli in ASD vs. TD [3=Low-level features]

With this third meta-analysis, we aim at further characterizing the low-level perceptual aspects potentially affecting the processing of BM in ASD vs. TD. Here, we focus only on studies that include both BM stimuli and non-BM scrambled stimuli. In other words, with this meta-analysis we inquire whether there are specific spatial or temporal features of the stimuli in motion that affect ASD and TD individuals differently.

The *I*^2^ coefficient indicates that the heterogeneity among the true effects is ∼58% (Test for Residual Heterogeneity: *QE*_(11)_ = 25.95, *p* =.007). Interestingly, results indicate that the moderator is significant, *QM*_(2)_ = 12.02, *p* =.003. To further investigate each level of the moderator we calculated separate meta-analyses for each moderator level. When considering only the studies with a BM task that includes a spatially scrambled stimulus, results indicated no significant difference in the performance of ASD vs. TD (Cohen’s *d* = 0.38, *SE* = 0.23, CI 95% = [-0.07, 0.83], *z* = 1.67, *p* =.10, Figure 8). The confidence interval includes 0, suggesting that we cannot exclude that the effect size for this subgroup of studies is null, and the heterogeneity is still moderate and significantly different from zero (*I*^2^ = 60%, *Q*_(6)_ = 14.84, *p* =.02).

**Figure 8.**
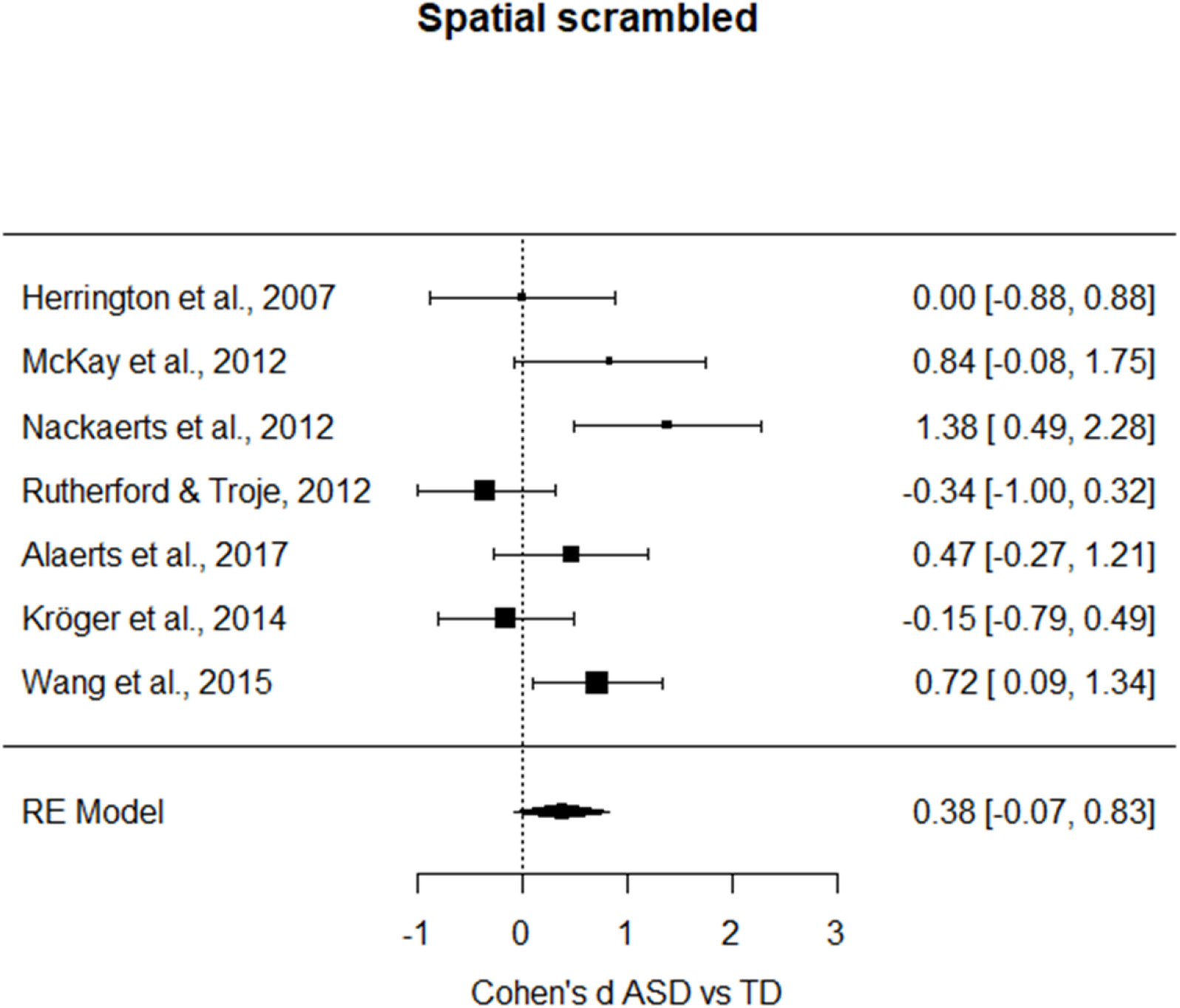
Forest plot of studies investigating the difference in BM performance between ASD and TD group that employed spatial scrambled stimuli (i.e. positive values indicate a better performance for the TD group as compared to the performance of the ASD group).

On the contrary, the model including the temporally scrambled stimulus showed a significant medium-large effect size (Cohen’s *d* = 0.67, *SE* = 0.21, CI 95% = [0.26, 1.08], *z* = 3.20, *p* =.001, Figure 9), and the included studies can be considered a homogenous group (*I*^2^ = 55%, *Q*_(5)_ = 11.11, p =.05). These differences in the processing of scrambled stimuli that we found were not confounded by the cognitive level, as it was shown by a one-way ANOVA on Cohen’s *d*s referring to cognitive level with ‘type of scramble’ as a factor (*F*_(1, 9)_ = 0.009, *p* =.93, *η_p_*^2^ = 0.001). Similarly, differences in the processing of scrambled stimuli were not confounded by the mean age of ASD groups (*F*_(1, 12)_ = 2.29, *p* = .16, *η_p_*^2^ = 0.16) nor by the mean age of TD groups (*F*_(1, 13)_ = 4.29, *p* =.06, *η_p_*^2^ = 0.25).

**Figure 9.**
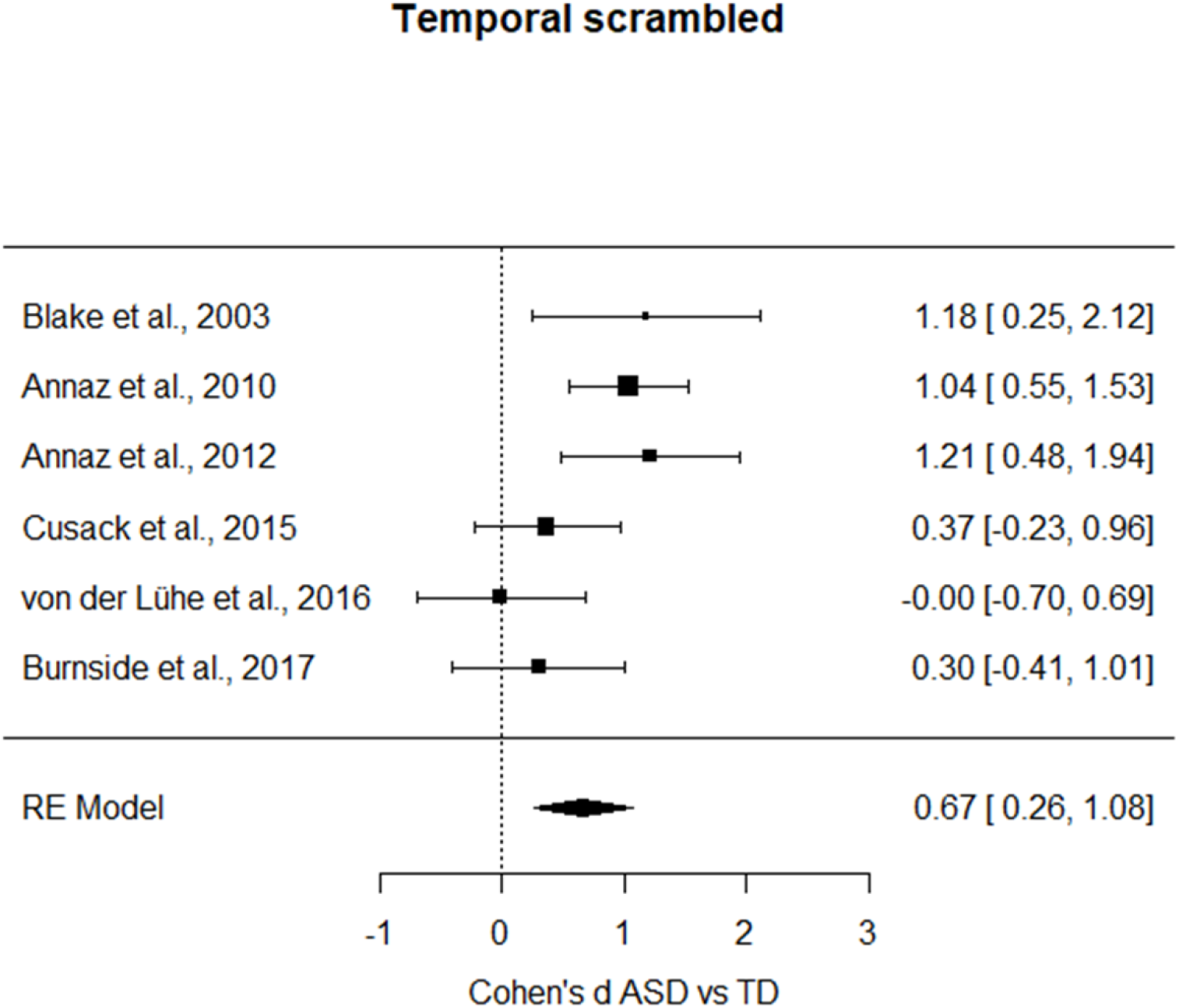
Forest plot of studies investigating the difference in BM performance between ASD and TD group that employed temporal scrambled stimuli (i.e. positive values indicate a better performance for the TD group as compared to the performance of the ASD group).

#### 3.5.4 Non-biological motion in ASD vs. TD [4=Non-BM]

From the included papers, we are here selecting only the studies including tasks that contrast BM stimuli with non-BM ones, to test whether the deficit is selective to BM stimuli or generalizes also to non-BM stimuli. Results indicate that no significant difference between the performance of individuals with ASD and TD is evident, (Cohen’s *d* = 0.26, *SE* = 0.29, CI 95% = [-0.31, 0.83], *z* = 0.89, *p* =.38, Figure 10). The total heterogeneity estimated with *I*^2^ is ∼75% (*Q*_(4)_ = 19.80, *p* =.0005).

**Figure 10.**
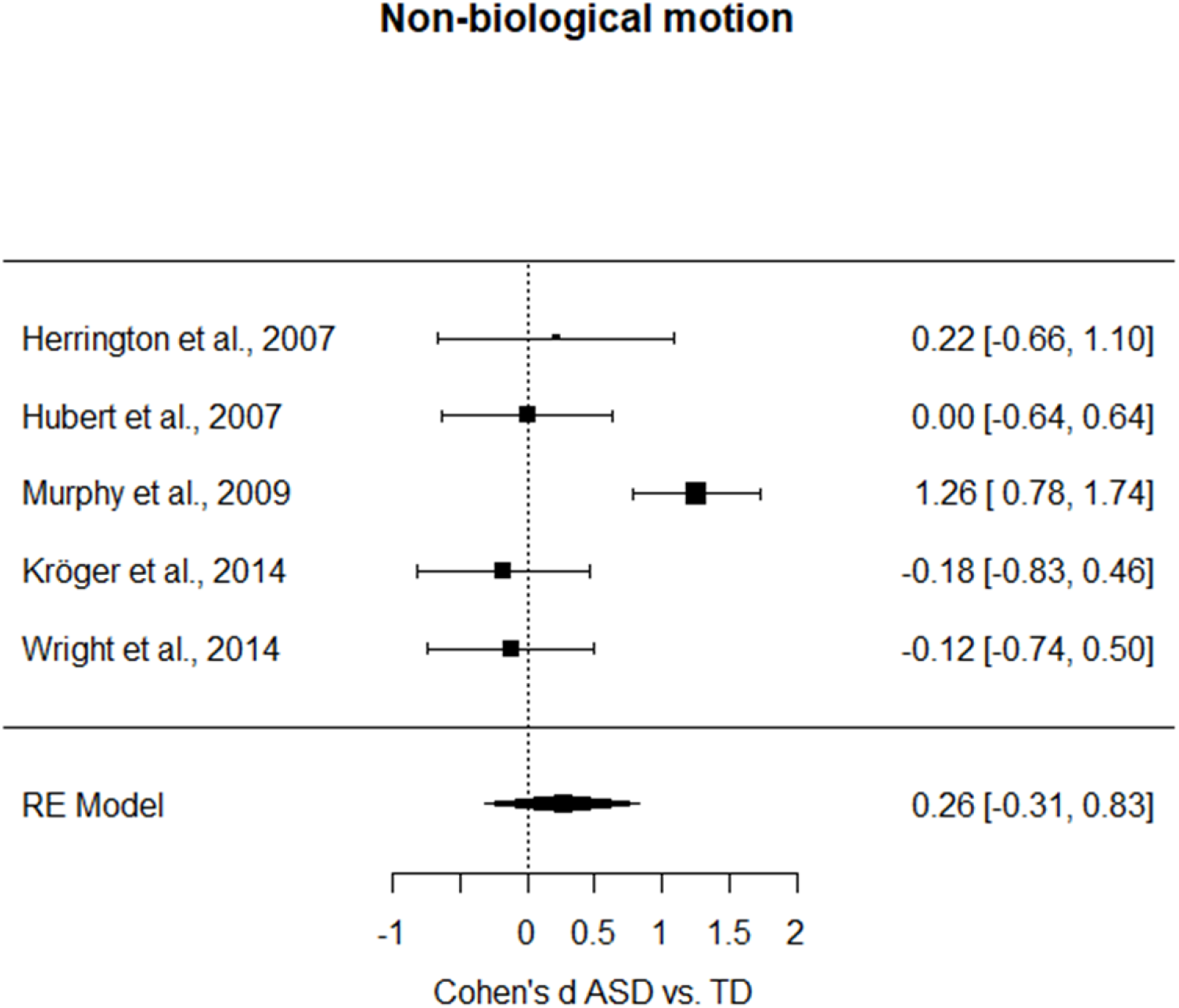
Forest plot of studies investigating the difference in the processing of non-BM between ASD and TD group (i.e. positive values indicate a better performance for the TD group as compared to the performance of the ASD group).

## 4. Discussion

In this article, we conducted the first meta-analytic investigation examining the abilities of individuals with ASD in the processing of BM stimuli. The results of the general meta-analysis [*1=All BM studies*] that included all the studies that matched our selection criteria showed a moderate (*d* = 0.57) deficit for individuals with ASD as compared to TD individuals (Table 2). This deficit was not influenced neither by the cognitive level nor by the mean age of participants and, moreover, it was not due to a publication bias. Nonetheless, we observed a high heterogeneity among studies (*I*² =79%), and this is an interesting aspect because it suggests that possible sources of variability are present in the group of studies included. We believe this represents a confirmation of the idea that the perception of BM is a complex ability, which cannot always be reduced to a monolithic process. Indeed, historically the BM literature has been characterized by remarkably heterogeneous experimental protocols. This can create ambiguity in transferring in operational terms the construct under investigation, which in turn can undermine reproducibility of studies, as previously proposed for example for the investigation of the “theory of mind” construct (Schaafsma et al., 2015). Thus, proven that the general meta-analysis points toward a significant impairment in BM perception in participants with ASD, we further developed our meta-analytic investigation in the direction of testing the possible factors that contribute to the high heterogeneity observed. Among these factors, we considered the distinct levels of processing defined in our three-level model (i.e., *first-order*, *direct* and *instrumental* BM processing), the type of manipulation used to create the comparison scrambled stimulus (spatial vs. temporal), and the selectivity of the impairment for biological as opposed to non-biological motion.

**Table 2.** Each row reports the main result for each of the four meta-analyses performed. Which meta-analysis, the subgroup, and the computed Cohen’s d are reported in the first, second and third column, respectively. The number of * represents the level of significance: *** p<.001; ** p<.01; * p<.05.

| META-ANALYSIS | SUBGROUP | EFFECT SIZE [CI] |
| --- | --- | --- |
| ***Biological motion in ASD vs. TD***<br>***[1=All BM studies]*** | / | 0.57 [0.30, 0.85] *** |
| ***Level of processing of biological motion in ASD vs. TD***<br>***[2=Level of processing]*** | *First-order* | 0.55 [0.02, 1.09] * |
|  | *Direct processing* | 0.40 [0.05, 0.74] * |
|  | *Instrumental processing* | 1.04 [0.68, 1.40] *** |
| ***Impact of low-level features of BM scrambled stimuli in ASD vs. TD***<br>***[3=Low-level features]*** | *Spatial Scrambled* | 0.38 [-0.07, 0.83] |
|  | *Temporal Scrambled* | 0.67 [0.26, 1.08] ** |
| ***Non-biological motion in ASD vs. TD***<br>***[4=Non-BM]*** | / | 0.26 [-0.31, 0.83] |

### 4.1 Is biological motion perception in ASD anomalous across all levels of processing?

A more precise analysis testing BM processing in ASD according to a three-level model that distinguished among *first-order*, *direct* and *instrumental* processing of BM confirmed that such a deficit was present independently of the specific level of information processing required in the BM task [*2=level of processing*]. Nonetheless, this moderator revealed a significant effect on the observed effect size of the impairment in BM processing, suggesting that the magnitude of this impairment depends on the level of processing involved in the specific type of task. Individuals with ASD showed the largest deficit as compared to TD participants when the experimental task required the *instrumental* recognition of the BM (*d* = 1.04), while this deficit was moderate (*d* = 0.55) for tasks requiring the *first-order* processing of BM and only small-to-moderate (*d* = 0.40) when tasks required the *direct* recognition of BM. It is worth noting that the category of *first-order* BM usually includes samples of younger children which performed preferential looking tasks. Moreover, for this category the deficit is moderate, but there is only a single study (Wright, Kelley, & Poulin-Dubois, 2016) in which we observed a negative effect size. This study however tested a preferential looking paradigm in children at the age of 6, and thus deviates from usual preferential looking experiments that usually include younger children or infants. Post-hoc tests revealed that *direct* and *instrumental* processing were significantly different in terms of effect size. The differences in the observed deficit as a function of the levels of BM processing were not ascribable to differences in cognitive level or mean age of ASD and TD groups across the three distinct levels of processing. The study of BM anomalies in ASD has so far been accumulating a conspicuous body of evidence, but often without putting the necessary effort in trying to understand the different sub-components required to efficiently process the distinct types of BM tasks. Taking advantage of the idea that BM can be processed at different levels of complexity, in this meta-analytic investigation we made a first attempt in this direction by distinguishing three levels in the BM processing. The most severe deficit in participants with ASD emerges when the recognition and categorization of a BM stimulus is serving a secondary purpose, such as when inferring someone else’s intentions, actions or emotional states; in other words, when correctly perceiving BM is a pre-requisite to disclose distinct and possibly more complex information. On the contrary, the least severe deficit is observable in participants with ASD when a more basic recognition of BM is tested, the level that we referred to as *direct* recognition of BM (e.g., BM vs. non-BM/scrambled stimulus; BM stimulus “A” vs. BM stimulus “B”, leftward vs. rightward moving stimulus, etc.). This less severe deficit could be due, at least in part, to compensatory mechanisms that can help participants with ASD to correctly process BM when the task is not excessively complex. Interestingly, a recent study in which morphed PLD stimuli were created by BM prototypical action, found that individuals with ASD showed intact abilities to recognize actions, but weakened adaptation to BM (van Boxtel, Dapretto, & Lu, 2016). This difference in the implicit processing of BM despite an intact explicit BM recognition suggests that individuals with ASD could develop compensatory mechanisms to perceive BM stimuli, which are more evident with increasing age. Accordingly, Rutherford and Troje (2012) have shown that, despite no group differences in sensitivity to BM or in the ability to identify the direction of motion were found between ASD and TD groups, only in the ASD group the IQ scores were significantly predictive of their performance. One possibility is that when participants with ASD are processing BM they use different strategies to compensate for their perceptual anomalies.

Overall, these results stress the need for a careful evaluation of what is at the core of BM processing and what are instead additional, secondary and higher-level information that can be extracted from a PLD depicting a stimulus moving in a biologically plausible fashion. Persevering with such an ambiguity would undermine the study of this construct as a putative endophenotype for ASD with negative clinical repercussions. For this reason, to assess how different low-level features can affect BM perception in ASD is pivotal.

### 4.2 What are the lower-level perceptual features linked to biological motion anomalies in ASD?

After assessing the role of different levels of BM processing in influencing the strength of the deficit that characterizes individuals with ASD, we assessed another fundamental aspect of BM that is the role of lower-level perceptual features [*3=Low-level features*]. Indeed, in many studies, BM stimuli are compared to scrambled stimuli obtained by varying the spatial or temporal properties of the PLD. Experiments using a *spatial* scrambled make use of comparison stimuli that are characterized by the displacement of all or part of the dots constituting the target PLD in a different spatial position but where each dot maintains the original temporal phase. Experiments using a *temporal* scrambled, instead, make use of comparison stimuli where each dot composing the target PLD remains in the same relative spatial position within the original configuration, but the motion of each of these dots is shown temporally out of phase. By testing the role of the type of scrambled stimuli used as a comparison, our aim was to inquire whether the deficits observable in individuals with ASD are dependent on the specific spatial or temporal properties of the target stimulus that are experimentally manipulated. We found that such a moderator had a significant influence in the effect size of the impaired BM processing in ASD. Studies employing a spatial scrambled as a comparison stimulus revealed no significant difference in the performance between individuals with ASD and TD controls. On the contrary, a significant impaired performance was evident for studies employing a temporal scrambled as a comparison stimulus, with a moderate-to-large effect size (*d* = 0.67). Although the sample size for each of this meta-analysis was limited (N=7 spatial; N=6 temporal), they furnish an initial interesting insight on the deficit underlying the perception of BM in ASD, which, we believe, deserves further investigation in future studies.

It is increasingly evident that simple spatio-temporal visual dynamics are essential building blocks for social behavior. A recent study in juvenile zebrafish showed that collective behavior, like shoaling, can be driven by simple specific motion cues that are constituted by black dots that mimic the precise zebrafish motion, while other naturalistic cues such as fish-like shape, pigmentation pattern, or non-visual sensory modalities are not required (Larsch & Baier, 2018). Anomalies in the processing stages connected to the tracking of low-level motion dynamics might be disruptive for detection, recognition and categorization of BM stimuli. Our results suggest that a candidate core mechanism playing a role in BM processing anomalies in ASD is linked to the temporal properties of moving dots embedded in a PLD. This is consistent with several experimental pieces of evidence and theoretical proposals suggesting that one of the core anomaly of individuals with ASD is related to temporal binding of uni- and -multi-sensory information (Brock, Brown, Boucher, & Rippon, 2002; Feldman et al., 2018; Wallace & Stevenson, 2014; Zhou et al., 2018), with accumulating evidence showing that this would be a common endophenotype also for other neurodevelopmental disabilities (Wallace & Stevenson, 2014; Zhou et al., 2018). Specifically, individuals with ASD have repeatedly shown an extended temporal binding window (TBW; an epoch of time within which stimuli from different sensory modalities are highly likely to be bound) which would bring to an imprecise temporal processing of sensory stimuli (Zhou et al., 2018). The majority of this evidence comes from audio-visual multisensory paradigms (for reviews see, Feldman et al., 2018; Wallace & Stevenson, 2014; Zhou et al., 2018), where researchers assess the temporal precision needed to perceive auditory and visual stimuli as synchronous. Nonetheless, initial evidence for temporal processing anomalies in unisensory perception has been documented in individuals with ASD in both uni-sensory auditory (Foss-Feig, Schauder, Key, Wallace, & Stone, 2017; Kwakye, Foss-Feig, Cascio, Stone, & Wallace, 2011b) and visual (Parks, 1965; J. I. R. Thompson et al., 2015) perception, suggesting a common underlying deficit in the fundamental coordination of brain networks that are responsible for the timing of sensory information processing (Burr & Morrone, 2006). It is also worth noting that the extension of TBWs has a high degree of malleability and they can shrink or expand as a function of the complexity and the low-level statistics of the incoming sensory information. For example, TBWs change as a function of spatial and temporal relationship among stimuli, such that for example information coming from different sensory modalities are integrated across a larger temporal window when they originate from the same spatial position (Zampini, Guest, Shore, & Spence, 2005) or when processing predictable/rhythmic stimuli as compared to irregular ones (Denison, Driver, & Ruff, 2012) (see also Casartelli, 2019). TBWs can shrink or expand also as a result of synchronization of brain rhythms (i.e. entrainment) (Ronconi, Busch, & Melcher, 2018; Ronconi & Melcher, 2017), or after few days of perceptual learning trainings (Powers, Hevey, & Wallace, 2012; Powers, Hillock, & Wallace, 2009; Stevenson, Wilson, Powers, & Wallace, 2013). Intriguingly, the hypothesis of a temporal processing anomaly as one of the key aspects responsible for BM deficits observed in ASD would be consistent with the involvement of the cerebellum and its bidirectional connectivity with the right posterior superior temporal sulcus (STS), which represent an important part of the network involved in the processing of BM (Jack, Keifer, & Pelphrey, 2017; Sokolov et al., 2012, 2014; Sokolov, Gharabaghi, Tatagiba, & Pavlova, 2010; Sokolov et al., 2018) as we will discuss in the next subsection.

### 4.3 Is the impairment observed in ASD specific to biological motion stimuli?

We conducted a meta-analysis of the subgroup of studies that tested performance in tasks with BM vs. non-BM stimuli [*4=Non-BM*]. The aim of this analysis was to assess whether the deficit of participants with ASD was selective for BM stimuli or generalized also to non-BM stimuli. We observed no evidence of impairments in the processing of non-BM stimuli in individuals with ASD; this suggests that the deficit is selective for BM stimuli and to their specific spatio-temporal pattern, whereas other types of complex stimuli in motion do not elicit a similar impairment. Among the non-BM tasks tested, we can find recognition/categorization of an object as a bike or a truck presented in PLD (Hubert et al., 2007; Wright et al., 2014) but also recognition/categorization of a scrambled version of the BM stimulus (Herrington et al., 2007; Kröger et al., 2014; Murphy et al., 2009). It is clear however that testing the scrambled version of BM stimuli is not ideal to claim that the deficit observed in individuals with ASD is specific to stimuli moving in a biological fashion. Consistently, there is evidence showing that when neurotypical adults cannot extract from a non-BM stimuli a form, non-BM stimuli themselves are more difficult to discriminate relative to BM stimuli. Specifically, these studies show that the presence of motion stimuli conveying specific forms nullifies the advantage in processing BM vs. non-BM stimuli. In other words, the presence of a form that can be extracted from non-BM stimuli seems to be a crucial aspect when comparing the recognition/categorization of BM and non-BM stimuli (Hiris, 2007). This would mean that BM is not generally easier to detect than equally-structured non-BM. To claim that individuals with ASD show a deficit which is specific to BM, the processing of non-BM stimuli should involve a similar extraction and interpretation of a form-from-motion. In the present meta-analysis there were only two studies that tested non-BM processing using tasks where the form was preserved (Hubert et al., 2007; Wright et al., 2014). These two studies, however, found no clear evidence of impairments in participants with ASD, independently of the nature of the moving stimuli (BM vs. non-BM) employed.

We conclude that from the evidence available at the time of the present meta-analysis, whether anomalies observed in individuals with ASD are specific to BM is still an open question. Thus, although interesting, the comparison between the processing of BM and non-BM stimuli needs a more careful investigation in future studies.

### 4.4 Neural and oscillatory correlates of BM processing in individuals with ASD

Looking at studies that investigate the neural correlates of BM processing in primates, neurotypical human, and in the ASD population, it seems to be consistent with the idea that BM is hardly conceivable as a monolithic process, given that the network of brain areas that are responsible for BM processing, although not yet completely understood, is highly complex and probably not unitary.

Neuroimaging studies in the typical population show that the activity of the posterior STS (pSTS) is specifically associated to BM stimuli presented as PLD. The pSTS involvement is often found to be lateralized in the right hemisphere (Brancucci, Lucci, Mazzatenta, & Tommasi, 2009; Grosbras, Beaton, & Eickhoff, 2012; Grossman et al., 2000; Pelphrey et al., 2003). While this region is activated for the observation of a wide range of human movements (Grosbras & Paus, 2006; Thompson, Hardee, Panayiotou, Crewther, & Puce, 2007), it has not been revealed to be involved in studies examining the perception of static bodies or body parts, suggesting that pSTS is involved specifically in extracting human body motion features rather than forms (Grosbras et al., 2012). Congruently, studies in non-human primates have shown the presence of neurons that selectively respond to various biological actions (e.g. walking, turning of the head, bending of the torso, moving of the arms, etc.; Jellema & Perrett, 2003). However, these neurons receive convergent information from the dorsal visual stream areas, middle temporal (MT) and medial superior temporal (MST), and also from the ventral stream area in the inferior temporal (IT) regions (Grosbras et al., 2012). The homologue of pSTS in the primate brain is also connected with the inferior parietal cortex (Seltzer & Pandya, 1994). A recent meta-analytic investigation of human brain regions activated in different types of BM processing (i.e. whole body, hand and face movements) confirmed common convergent activations independent of the specific motion category in the right pSTS region. However, the pattern of activations included also bilateral regions at the junction between middle temporal and lateral occipital gyri (Grosbras et al., 2012).

Other evidence in favor of a highly complex network of brain regions supporting the processing of BM that goes well beyond pSTS comes from functional connectivity studies. For example, a study by Sokolov et al. (2012) that used fMRI to test healthy participants during a BM task, in combination with functional connectivity analysis and dynamic causal modeling, showed that the left lateral cerebellum (lobules Crus I and VIIB) exhibited increased activation and connectivity with right pSTS during the task. More recently, Sokolov et al. (2018) used novel dynamic causal modeling informed by probabilistic tractography to highlight that the network for BM processing is organized in a parallel rather than hierarchical manner. Specifically, the authors showed that although the right pSTS serves as an integrator within temporal areas, this area does not appear to be the main gate in the functional integration of the activity of the occipito-temporal and frontal regions. Indeed, both fusiform gyrus and middle temporal areas were also connected to the right inferior frontal gyrus and insula during processing of BM stimuli, indicating multiple parallel pathways. BM-specific loops of effective connectivity were found also between the left lateral cerebellar lobule Crus I and right pSTS, as well as between the left Crus I and right insula. According to their conclusions, this parallel processing that is dependent on the activity of different cortico-cortical and cortico-cerebellar pathways may help to explain why BM processing is rather resilient to focal brain damage; in contrast, it has been hypothesized to be anomalous in neuropsychiatric conditions with distributed network alterations (e.g., ASD), as widely debated in our work.

In addition, it has been suggested that the cerebellum uses a common computational algorithm not only upon motor but also non-motor functions, such as perceptual ones (D’Angelo & Casali, 2012; Knolle, Schröger, & Kotz, 2013). The idea that the cerebellum may play a critical role as internal “timing device” for motor and non-motor functions has been widely explored in the literature (Bareš et al., 2018). Notably, this idea may fit with the evidence indicating that the cerebellum is implicated in the pathophysiology of ASD (Fatemi et al., 2012; Wang et al., 2014) and may also fit with our present results showing a specific role for temporal scrambled stimuli in explaining the anomalies in BM processing observed in individuals with ASD.

Although it is not within the scope of the present article to provide an extensive review of the neuroimaging literature that investigates the neural correlates of impaired BM processing in ASD, it is interesting to notice that consistent report in the literature show that during the processing of BM stimuli individuals with ASD show hypoactivation in the pSTS (Ahmed & Vander Wyk, 2013; Freitag et al., 2008; Herrington, Nymberg, & Schultz, 2011; Kaiser et al., 2010) and in regions of the prefrontal cortex (Kaiser et al., 2010; Koldewyn et al., 2011a). However, only few studies so far have tested the effective connectivity among the key regions of the cortical and subcortical (i.e. cerebellum) network responsible of the processing of BM (Alaerts, Swinnen, & Wenderoth, 2017; Jack et al., 2017; McKay et al., 2012). These studies suggest that a defective temporo-parietal (McKay et al., 2012) and cerebellar-pSTS connectivity (Jack et al., 2017) could contribute to the anomalies in the processing of BM reported in ASD and reviewed in the present meta-analysis. Although these studies provide initial important evidence, the results of the present meta-analysis suggest that distinct levels of BM processing can be differentially affected in participants with ASD, and that the manipulation of low-level perceptual features may result in markedly distinctive behavioral outcomes. These aspects should be taken into account in future neuroimaging studies that will address the neural correlates of the anomalies in the processing of BM in ASD. More generally, even from a neural perspective, BM should not be considered as a monolithic process, but as constituted by different operations, that are likely supported by different parallel neural routes.

### 4.5 Limitations and future directions

Although not conclusive, we believe that the model and the cumulative findings reported in the present meta-analysis move the conceptualization of BM anomalies in ASD a step forward. To improve the explanatory power of this and other models proposed in the field of BM, we call for future studies to provide appropriate quantitative data to be summarized with metanalytic approaches. Specifically, we deem relevant to investigate whether the potential anomalies in BM processing in ASD are extended to non-BM stimuli, and whether they are selectively tied to a temporal binding deficit. To this end, testing multiple BM and non-BM stimuli (e.g., BM, non-BM, spatial scrambled, temporal scrambled) within the same ASD sample would be of paramount importance (please refer to meta-analysis 3 = *Low-level features* and 4 = *Non-BM*).

Although the current high variability in experimental questions prevented us from conducting a meta-analysis on more aspects of BM processing in ASD, we find extremely interesting to produce more evidence on the specificity of the ASD impairment in BM vs. other motion tasks (e.g., coherent dot motion or the form-from motion tasks). Indeed, in two of the studies included in our meta-analysis a coherent motion task was used as a control task (Atkinson, 2009; Koldewyn, Whitney, & Rivera, 2011). Results indicated that in the ASD group the performance to BM and coherent motion tasks are positively correlated. Future studies on contrasting in the same sample the performance to BM, coherent motion and form-from motion tasks will aid the clarification of whether the deficit in BM processing showed by ASD individuals is specific to BM or instead is common to other complex motion tasks. In accordance with the idea of general perceptual anomalies in ASD that are not specific to BM, a recent study (Falck-Ytter et al., 2018) has shown that infants that are less attracted to multisensory BM stimuli were diagnosed with ASD at 2 years of age. Future works in this direction will clarify how anomalies in BM processing in ASD, if any, should be appropriately conceived for clinical aims.

## 5. Conclusions

Taken together, this work represents to our knowledge the first meta-analytic investigation that assesses in a detailed manner the putative impairment in the processing of BM in individuals with ASD. Despite some of our analyses are probably influenced by the limited sample of studies available and by the high heterogeneity of participants’ characteristics, we made an effort toward interpreting the multifaceted aspects of the vast literature that uses BM as a proxy for social perception deficits in ASD. While on the one hand a significant impairment in BM processing emerged in individuals with ASD, on the other hand the idea that this deficit is selective for BM and does not extend to non-BM stimuli is, at present, not properly supported by the literature. Moreover, we observed a high heterogeneity due to the different experimental protocols employed. A major source of heterogeneity can be individuated in the distinct levels (*first-order, direct, instrumental*) of BM processing, given that the most severe deficit in participants with ASD emerges when processing of a BM stimulus is serving a secondary purpose (e.g. inferring other’s intentionality, action or emotional state). How more severe difficulties reported in the *instrumental* recognition of BM may represent just a byproduct of more basic anomalies should be clarified in future studies. Finally, an initial result points toward the importance of lower-level temporal perceptual features in determining the deficit in the perception of BM in ASD.

## Acknowledgements

This work was supported by the award fellowship granted by SISSA to V.P., and by a grant from the Italian Ministry of Health (RC 2016-2018) to L.C.. The funders did not participate in the conception and development of this work. We additionally want to warmly thank the authors of the papers included in the meta-analysis who sent us the data we required along the process.

## Authors’ contribution

A.F., V.P., L.C. and L.Ro. have conceived the idea of the paper. A.F. and L.Ra. executed the literature search, coded the papers and calculated the effect sizes, independently. A.F. and M.V. defined the analytical protocol and performed the analyses. A.F., V.P., L.C. and L.Ro. drafted the article, and all authors critically revised it. All authors approved the final version of this article.

## Footnotes

^1^ For very small sample sizes (i.e., *n*_1_ < 10 and *n*_2_ < 10), Cohen’s *d* overestimates both the effect size and its variance, therefore the use of Hedges’ *g* is sometimes preferred over the use of Cohen’s *d* as a measure of the effect size (see Lakens, 2013). However, as reported by Borenstein et al. 2009, p. 28, for sample sizes larger than 10 the difference between Cohen’s *d* and Hedges’ g is trivial, which led us to favor the use of the easily interpretable and very well-known Cohen’s *d*.

^2^Note that, when a large number of studies include a large number of multiple measures, the robust variance estimation method (Tanner-Smith et al., 2016) is sometimes used to obtain unbiased estimates of the average Cohen’s *d* and of its variance. However, because in the current context the number of studies including multiple effect size measures was relatively small (i.e., 10/27 in meta-analysis 1), and because there were no more than three effect size measures for each of these studies (more often there were two), we deemed it preferable using a simpler random-effects model with appropriate methods for aggregating effect sizes from the same study (Borenstein et al., 2009).

